# Late-in-life treadmill-training rejuvenates autophagy, protein aggregate clearance, and function in mouse hearts

**DOI:** 10.1101/2021.05.27.446031

**Authors:** Jae Min Cho, Kellsey Ly, Caroline Ramous, Lauren Thompson, Michele Hansen, Maria Sara de Lima Coutinho Mattera, Karla Maria Pires, Maroua Ferhat, Kevin J Whitehead, Kandis Carter, Márcio Buffolo, Rajeshwary Ghosh, Seul-Ki Park, Sihem Boudina, J David Symons

## Abstract

There is evidence for a progressive decline of protein quality control mechanisms during the process of cardiac aging. This enables the accumulation of protein aggregates and damaged organelles that contribute to age-associated cardiac dysfunction. Macroautophagy (referred to as autophagy) is the process by which post-mitotic cells such as cardiomyocytes clear defective proteins and organelles. We hypothesized that late-in-life exercise training improves autophagy, protein aggregate clearance, and function that is otherwise dysregulated in hearts from old vs adult mice. As expected, 24-month old male C57BL/6J mice (old) exhibited : (i) repressed autophagosome formation and protein aggregate accumulation in the heart; (ii) systolic and diastolic dysfunction; and (iii) reduced exercise capacity, vs. 8-month old (adult) mice (all p< .05). Separate cohorts of 21 month old mice completed a 3-month progressive resistance treadmill-running program (old-ETR) that improved (all < .05) : (i) body composition; (ii) exercise capacity; and (iii) soleus muscle citrate synthase activity, vs. age-matched mice that did not train (old-SED). Importantly, (iv) protein expression of autophagy markers indicated trafficking of the autophagosome to the lysosome increased, (v) protein aggregate clearance improved, and (vi) overall function was enhanced (all p<0.05), in hearts from old-ETR vs. old- SED mice. Dietary maneuvers and pharmacological interventions shown to elevate basal autophagy are reported to mitigate / reverse age-associated cardiac dysfunction. Here we show the first evidence that a physiological intervention initiated late-in-life improves autophagic flux, protein aggregate clearance, and overall function in mouse hearts.

## 1 INTRODUCTION

The incidence of cardiovascular disease (CVD) is ∼20%, ∼50%, ∼80%, and ∼90% in individuals 18-44, 45-64, 65-79, and 80+ years of age, respectively (Benjamin et al., 2019). Treating cardiovascular complications associated with the aging demographic creates an enormous economic challenge to the healthcare system in particular, and society in general. New therapeutic targets need to be identified so that practical intervention strategies can be designed, optimized, and implemented. We sought to determine whether myocardial autophagy can be influenced positively by a well-accepted lifestyle intervention strategy (i.e., regular physical activity) in a preclinical model of primary aging.

Protein aggregates accumulate and organelles become damaged and / or dysfunctional during the process of aging. A progressive loss of the cellular quality control mechanism autophagy contributes importantly in many organs to this age-associated proteotoxicity and the subsequent decline in cellular function (Cuervo & Dice, 2000; Donati, Recchia, Cavallini, & Bergamini, 2008; Koga, Kaushik, & Cuervo, 2011; Rubinsztein, Marino, & Kroemer, 2011; Vittorini et al., 1999). Post-mitotic cells with limited proliferative capacity such as cardiac myocytes are particularly reliant upon autophagy to maintain proteostasis and thereby preserve myocardial function during aging (Rubinsztein et al., 2011; Terman & Brunk, 2005). In support of this, age-related cardiomyopathy is recapitulated in adult mice by cardiac specific Atg5 deletion (Taneike et al., 2010) and mTORC1 activation (Li et al., 2017; Taneike et al., 2016; Zhou et al., 2013), whereas desmin-related cardiomyopathy, characterized by the accumulation of cytotoxic misfolded proteins, is prevented by cardiac-selective Atg7 overexpression (Bhuiyan et al., 2013).

Most literature indicates that primary aging precipitates myocardial dysfunction in C57BL/6J mice (Dai & Rabinovitch, 2009; Dai, Rabinovitch, & Ungvari, 2012; Dai et al., 2009), but comparisons of cardiac autophagy between older and adult mice have not yielded consistent findings (Boyle et al., 2011; Hua et al., 2011; Inuzuka et al., 2009; Li et al., 2017; L. Ma et al., 2017; Marin-Aguilar et al., 2020; Peng et al., 2013; Ren et al., 2017; Taneike et al., 2010; Wang et al., 2018; Wu et al., 2016; Zhou et al., 2017; Zhou et al., 2013). Inconsistencies likely arise from conclusions being based solely upon steady-state measures of autophagy markers including MAP1LC3/LC3 (microtubule associated protein 1 light chain 3) and SQSTM1/p62 (Klionsky et al., 2016; Mizushima, Yoshimori, & Levine, 2010). However, because autophagy is a highly dynamic process it is best practice to pharmacologically block the turnover of these proteins to most accurately quantify the scale of autophagosome formation i.e., autophagic flux. A review of the literature reveals this methodological approach has been implemented once in aged mice (Wu et al., 2016), and once in cardiomyocytes isolated from older rats (L. Ma et al., 2017). Both studies provided support for an age-associated repression of cardiac autophagic flux. Here we substantiated these observations and additionally showed accrual of ubiquitinated proteins and protein aggregates in the myocardium, cardiac dysfunction, and reduced exercise capacity, in 24- mo vs. 8-mo old animals. These findings allowed us to test our primary hypothesis.

A growing area of research inquiry is whether upregulating the process of autophagy has therapeutic benefit. For example, late-in-life interventions known to increase autophagy such as supplementation with the natural polyamine spermidine (Eisenberg et al., 2016), caloric restriction (Sheng et al., 2017), and mTORC1 inhibition using rapamycin (Flynn et al., 2013), lessen age-associated cardiac dysfunction in C57BL/6J mice. An alternative or complementary approach with potential to improve myocardial autophagy and attenuate the aging associated decline in cardiac function is dynamic exercise. In this regard, He et al., observed that an acute bout of treadmill-running elevates protein indexes of autophagy in murine hearts (He et al., 2012), and Bhuiyan et al. reported that autophagy, protein clearance, and function improved in myocardium from mice with desmin-related cardiomyopathy that did vs. did not have long-term access (i.e., 6-mo) to wheel-running (Bhuiyan et al., 2013). The potential for a physiological maneuver i.e., late-in-life exercise-training, to rejuvenate myocardial autophagy has not been investigated. Here we present what we believe to be the first report that exercise capacity, autophagic flux, protein clearance, redox balance, and cardiac function improve in hearts from older mice that complete a progressive, resistance treadmill-running program vs. animals that do not train.

## 2 RESULTS

### 2.1 Myocardium from older vs. adult mice displays repressed autophagic flux

The impact of aging on cardiac autophagy in pre-clinical murine models is not uniform (Boyle et al., 2011; Hua et al., 2011; Inuzuka et al., 2009; Li et al., 2017; L. Ma et al., 2017; Marin-Aguilar et al., 2020; Peng et al., 2013; Ren et al., 2017; Taneike et al., 2010; Wang et al., 2018; Wu et al., 2016; Zhou et al., 2017; Zhou et al., 2013). Few studies have assessed the influence of aging on different steps involved in the process of myocardial autophagy (L. Ma et al., 2017; Wu et al., 2016). To address both issues, adult and older male C57BL/6J mice completed TD-NMR analyses to determine body composition. This was required so that the lysosomal acidification inhibitor chloroquine (CQ) could be administered using a dose based on lean body mass (Glick, Barth, & Macleod, 2010; Gottlieb, Andres, Sin, & Taylor, 2015; Pires et al., 2017). A schematic of our procedures is shown in Figure 1a. Twenty-four h after TD-NMR, CQ (75 mg IP / g lean body mass) or vehicle-control (phosphate-buffered saline; VEH) (Gottlieb et al., 2015) was administered to both groups. After 4 h hearts were collected from isoflurane anesthetized mice. Based on available literature, we hypothesized that autophagic flux would be impaired in hearts from older vs. adult mice, and that this would associate with accrual of ubiquitinated proteins and heightened oxidant stress in the older hearts. Representative images (Figure 1b) and mean data indicate protein expression of LC3I:GAPDH, LC3II:GAPDH, and p62:GAPDH was higher (Figure 1c,d,f; p < .05), and Atg3:GAPDH was lower (Figure S1a,b; p < .05), in myocardial homogenates from old-VEH vs. adult-VEH mice (histogram 3 vs 1), whereas LC3II:LC3I, Atg5:GAPDH, and Atg7:GAPDH (Figures 1e and S1a,c,d) were similar between groups. Atg3 mRNA expression was lower (p < .05) in myocardium from old-VEH (0.87 ± 0.02) vs. adult-VEH (1.00 ± 0.04) mice. These findings were concurrent with accrual (p < .05) of ubiquitinated proteins (Figure S2a,b), elevated (p < .05) 4-hydroxy-2-nonenal (4-HNE; Figure S2c,d) and superoxide dismutase (SOD) 2 (Figure S2e), and reduced (p < .05) catalase (Figure S2e).

**Figure 1.**
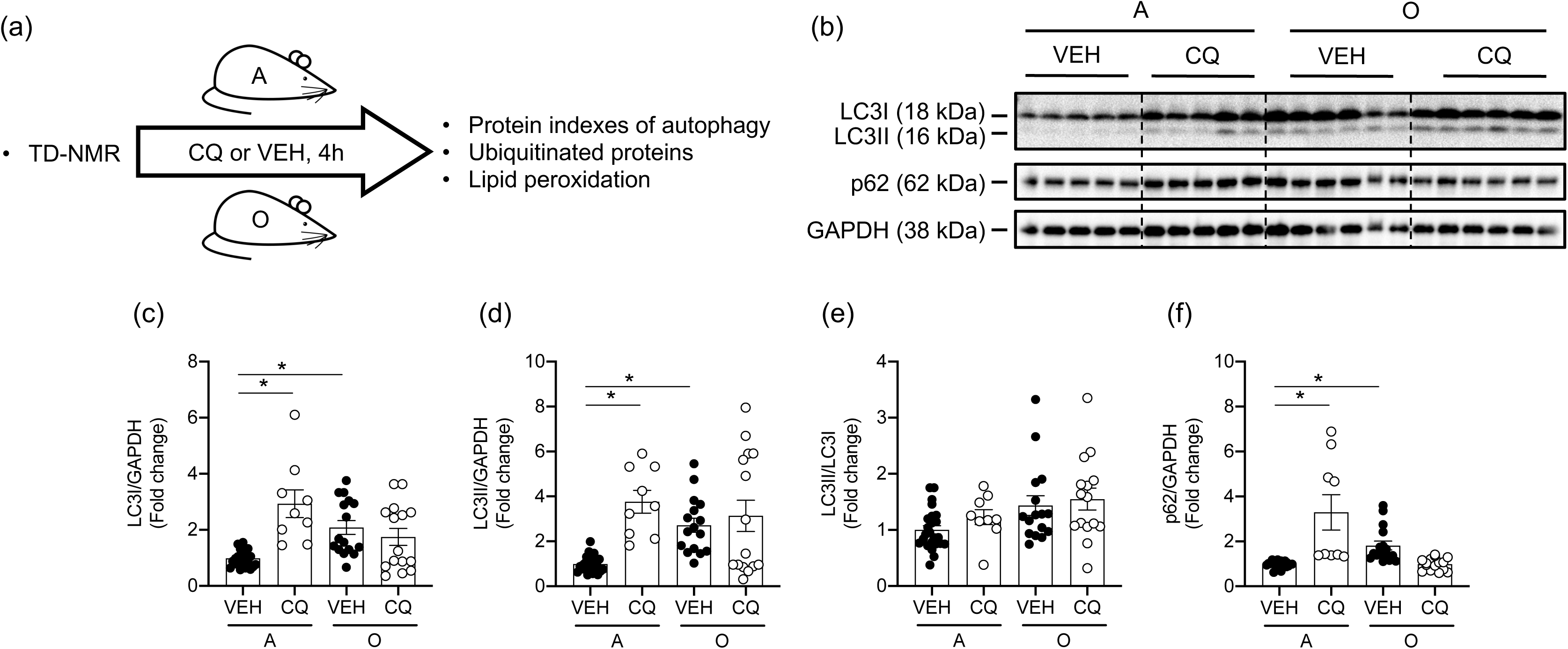
*Autophagic flux is robust in hearts from adult but not older mice.* (a) Body composition was assessed using TD-NMR in adult (A; 8-mo) and older (O; 24-mo) mice. Vehicle (VEH; phosphate buffered saline) or chloroquine (CQ; 75 mg CQ / g of lean body mass) was administered IP to A and O mice. Four h later mice segments of myocardium were obtained and prepared for immunoblotting. Representative images (b) and mean data ± standard error are shown (c-f). LC3-I, LC3-II, and p62 normalized to GAPDH were greater in myocardium from O vs. A mice treated with vehicle, whereas LC3-II:LC3-I was similar between groups. In A mice, LC3-I, LC3-II, and p62 normalized to GAPDH increased in CQ vs. VEH treated cohorts, whereas these endpoints were similar in O mice treated with CQ and VEH. These findings demonstrate that autophagic flux is robust in A but not O mice. For panels (c-f), n=9-24, *p<0.05 vs A mice. Data are expressed as fold change relative to values obtained from A mice.

It is not conclusive whether aging impairs autophagosome formation, trafficking of the autophagosome to the lysosome and / or lysosomal degradation of the autophagosome in the heart. Representative images (Figure 1b) and mean data indicate protein expression of LC3I:GAPDH, LC3II:GAPDH, and p62:GAPDH was higher (Figure 1c,d,f; p < .05) in myocardial homogenates from adult-CQ vs. adult-VEH mice (histogram 2 vs. 1). Elevated (p < .05) LC3-II:GAPDH and p62:GAPDH in old-VEH vs. adult-VEH mice (histogram 3 vs. 1) did not increase further in older mice treated with CQ (histogram 3 vs. 4; Figure 1b-f), supporting the hypothesis that cardiac autophagic flux is impaired by aging at the step of autophagosome clearance.

### 2.2 Myocardial function is impaired in older vs. adult mice

While some discrepancies exist, (Barouch et al., 2003; H. Ma et al., 2010) the balance of available literature indicates that primary aging precipitates myocardial dysfunction in C57BL/6J mice (Dai & Rabinovitch, 2009; Dai et al., 2012; Dai et al., 2009). Based on the majority of findings to date, we hypothesized that aging-associated myocardial dysfunction exists and a schematic of our procedures is shown in Figure 2a. Compared to adult mice, older animals displayed greater left ventricular (LV) mass / tibia length (Figure 2b; Table S1), lower EF, FS, CO (Figure 2c,d,f), and higher LVIDs (Figure S3a; all p < .05), indicating LV hypertrophy and systolic dysfunction exist in hearts from aged animals. Further, older mice exhibited diastolic dysfunction e.g., lower mitral valve flow E wave velocity (MVE) and A wave velocity (MVA) (Figure 2g,h; p < .05), together with a trend toward an elevated E/A ratio (Figure 2i; p=0.07). The myocardial performance index (MPI), an indicator of overall LV function, is calculated by the (ICT+IRT) / ET, where ICT is isovolumetric contraction time, IRT is isovolumetric relaxation time, and ET is ejection time (Goroshi & Chand, 2016; Tei et al., 1995). MPI was elevated (i.e., function became worse) in hearts from older vs. adult mice (Figure 2k; p < .05). Of note, the accrual of p62:GAPDH (indicating repressed autophagy) associated (p < .001) with increased MPI (worsening of cardiac function; Figure 2l).

**Figure 2.**
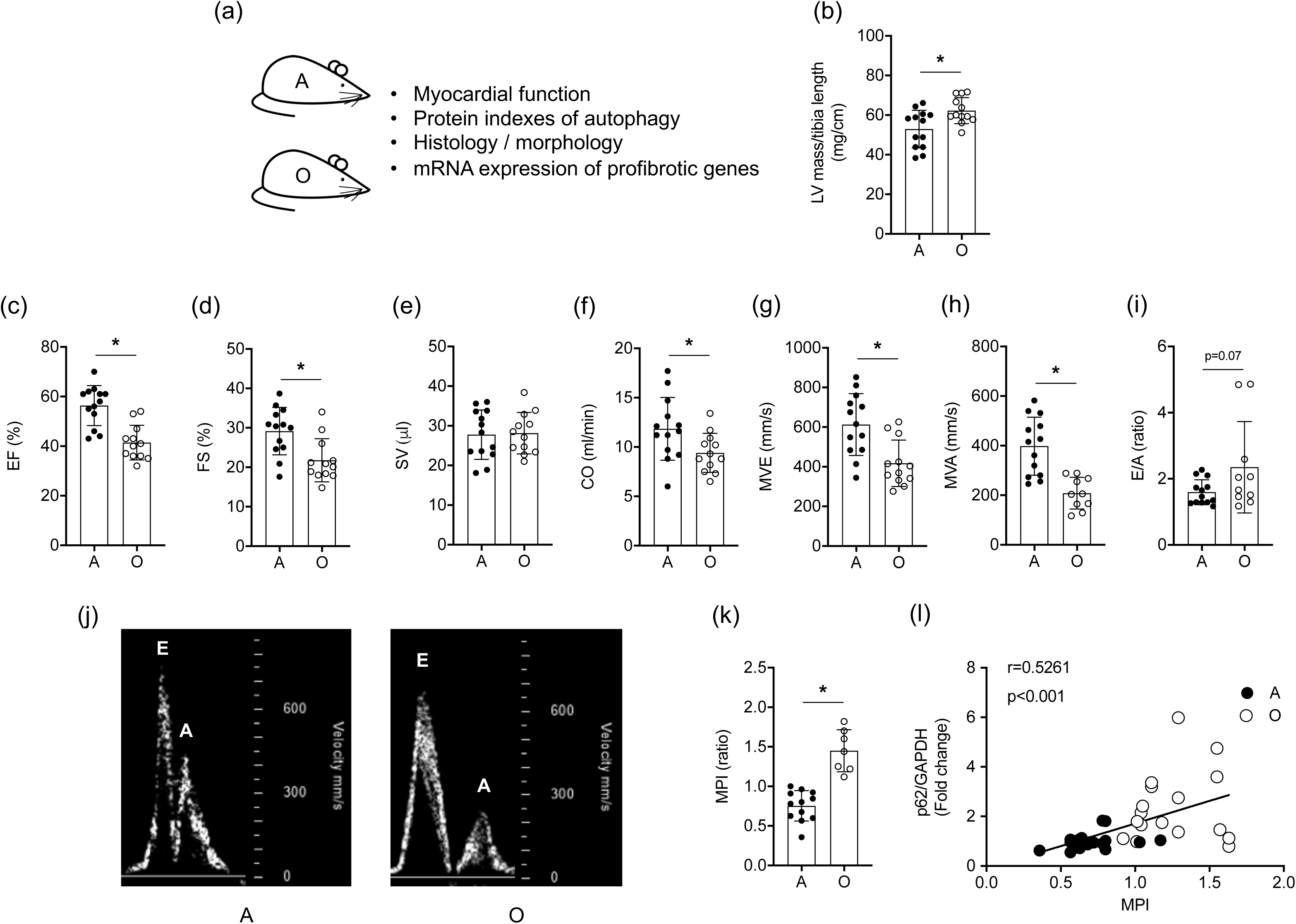
*Myocardial function is impaired in older vs. adult mice.* (a) Transthoracic echocardiography was performed on adult (A; 8-mo) and older (O; 24-mo) mice. Mean data ± standard error are shown (b-i, k). Left ventricular (LV) mass normalized to tibial length was greater in O vs. A animals. Ejection fraction (EF, %; c), fractional shortening (FS, %; d), cardiac output (CO, ml/min; f), passive diastolic filling (MVE, mm/s; g), and active diastolic filling (MVA, mm/s; h) were impaired in O vs. A mice, whereas stroke volume (SV, μl; e) and the E/A ratio (i) were not different between groups. These data, together with our observation that the myocardial performance index (MPI, k) is greater in hearts from O vs. A mice, substantiate that, on balance, systolic and diastolic dysfunction exists in O vs. A mice. (j) Representative images of blood flow velocity obtained during the assessment of MVE and MVA from both groups. (l) The correlation between protein expression of p62/GAPDH and MPI was strong in hearts from A and O mice. For (b-i, k) n=12-13; for (l) n=16-17. *p<0.05 vs A.

While type 1 collagen staining indicated increased fibrosis in hearts from older vs. adult mice, cardiomyocyte area was not affected (Figure S4a-c). Of the profibrotic genes that were assessed, mRNA expression of fibrillin (Fbn) 1 and transforming growth factor-β (Tgfb) 2 were elevated (p < .05), and a trend existed for increased connective tissue growth factor (Ctgf; p=0.07) in hearts from older vs. adult mice, whereas Fbn2 and Tgfb1 were not different between groups (Figure S4d; Table S2).

### 2.3 Exercise-training is efficacious in adult and older mice

Our findings of repressed autophagic flux (Figures 1 and S1), ubiquitinated protein accrual and heightened oxidant stress (Figure S2), and impaired function (Figure 2), in hearts from older vs. adult mice were anticipated. However, it was necessary for us to substantiate these observations to determine whether a physiological intervention i.e., exercise-training, lessens these age-associated disruptions. To achieve this, adult mice did (adult-ETR) or did not (adult-SED) complete progressive resistance treadmill-training from 5 to 8 months of age. Likewise, older mice did (old-ETR) or did not (old-SED) train from 21-24 months of age. A schematic is shown in Figure 3a. As expected, fat mass and body mass were greater (Figure S5a,b; p < .05), whereas total workload capacity and soleus muscle citrate synthase (CS) enzyme activity were less (Figure S5e,f; p < .05), in old-SED vs. adult-SED (histogram 3 vs. 1) mice (Table S1). Evidence for an exercise-training effect included : (i) reduced fat mass (Figure S5a), and increased total workload capacity (Figure S5e) and CS activity (Figure S5f), in adult (histogram 2 vs. 1) and older (histogram 4 vs. 3) mice (p < .05 for all; Figure S5).

**Figure 3.**
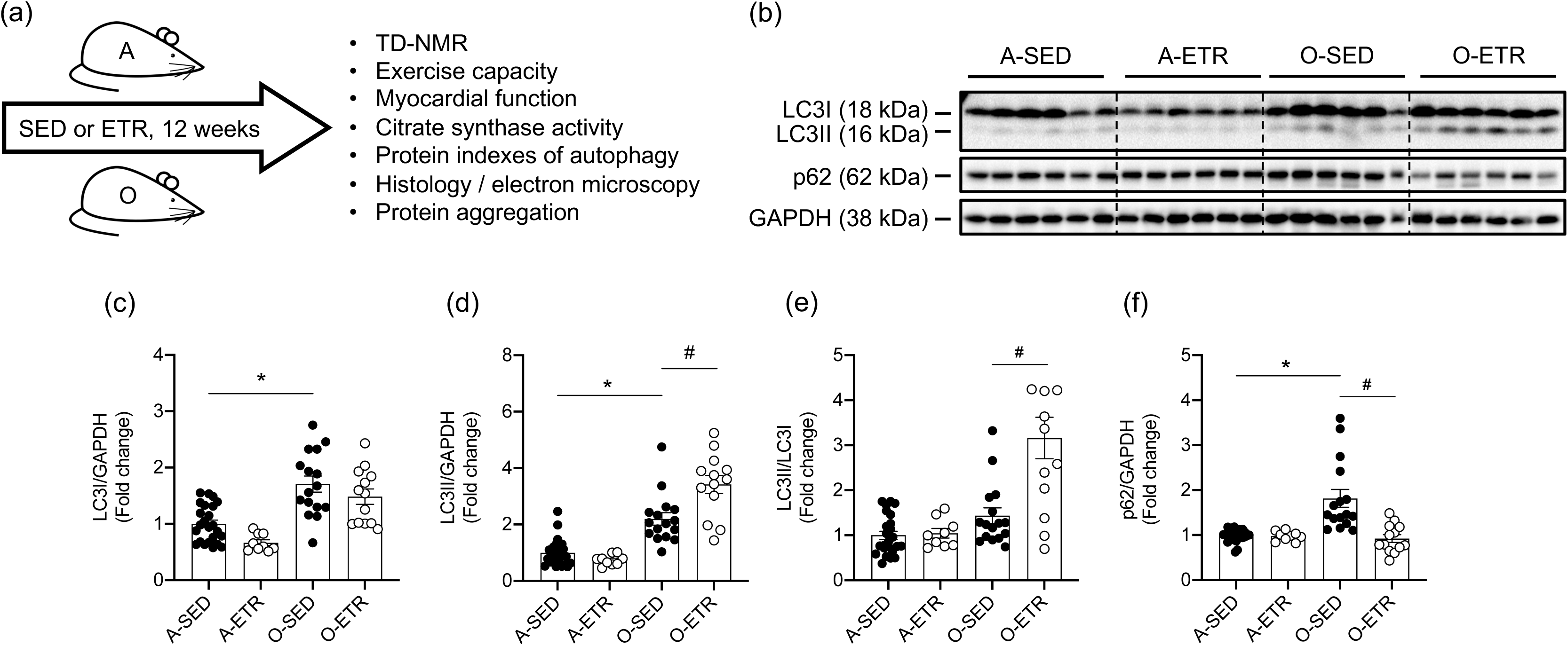
*Steady-state autophagy is improved by exercise-training in hearts from older but not adult mice.* (a) Adult (A, 5 mo) and older (O, 21 mo) mice did (ETR) or did not (SED) complete 12-weeks of treadmill-running. (b) At least 24 h following the last exercise bout, A (8 mo) and O (24 mo) mice were anesthetized and segments of myocardium were prepared for immunoblotting. Representative images (b) and mean data ± standard error are shown (c-f). LC3-I, LC3-II, and p62 normalized to GAPDH were greater in myocardium from O-SED vs. A-SED mice, whereas LC3-II:LC3-I was similar between groups. This pattern is identical to the one shown for O and A mice in Figure 1. While no differences existed between A-SED and A-ETR mice, LC3-II (d) normalized to GAPDH and LC3II/LC3I were increased in O-ETR vs. O- SED. p62 normalized to GAPDH decreased in O-ETR vs. O-SED animals. For panels (c-f), n=9-24, *p<0.05 vs A-SED; #p<0.05 vs O-SED. Data are expressed as fold change relative to values obtained from A-SED mice.

### 2.4 Exercise training improves autophagic flux in myocardium from older mice

Substantiating findings shown in Figure 1 that steady state autophagy is impaired in hearts from older vs. adult mice, representative images (Figure 3b) and mean data indicate protein expression of LC3I:GAPDH, LC3II:GAPDH, and p62:GAPDH was higher (Figure 3c,d,f; p < .05), and Atg3:GAPDH was lower (p < .05; Figure S6a,b), in myocardial homogenates from older-SED vs. adult-SED mice (histogram 3 vs. 1). Regarding exercise-training, indexes of cardiac autophagy were similar between adult-ETR and adult-SED mice (histogram 2 vs. 1; Figures 3 and S6). In contrast, LC3I:GAPDH and p62:GAPDH were less (p < .05; Figure 3c,f), and Atg3 protein expression was greater (p < .05; Figure S6a,b), in hearts from old-ETR vs. old-SED mice (histogram 4 vs. 3). Atg3 mRNA expression was increased (p < .05) in myocardium from old-ETR (1.45 ± 0.11) vs old-SED (1.00 ± 0.17) mice.

Next we discerned whether exercise-training improves the aging-associated repression of autophagic flux observed in Figure 1. A schematic of our approach is shown in Figure 4a. Old-SED and O-ETR mice completed TD-NMR so that CQ could be administered based on lean body mass. Twenty-four h later older-SED and older-ETR mice were treated with CQ or VEH. These experiments were not performed using adult mice because exercise training did not influence steady state indexes of autophagy in that cohort of animals (Figure 3). Substantiating results shown in Figure 3, p62:GAPDH protein degradation improved (p < .05) in old-ETR vs. old-SED mice (Figure 4, histogram 3 vs. 1). Supporting findings shown in Figure 1, indexes of autophagy assessed in old-SED mice did not respond to CQ treatment (Figure 4, histogram 2 vs. 1). Notably, p62:GAPDH accrual (Figure 4f) and Atg3:GAPDH accumulation (Figure S7a,b; p < .05) occurred in the presence vs. the absence of CQ in old-ETR animals (histogram 4 vs. 3). These findings provide support that exercise-training improves autophagic flux in hearts from older mice.

**Figure 4.**
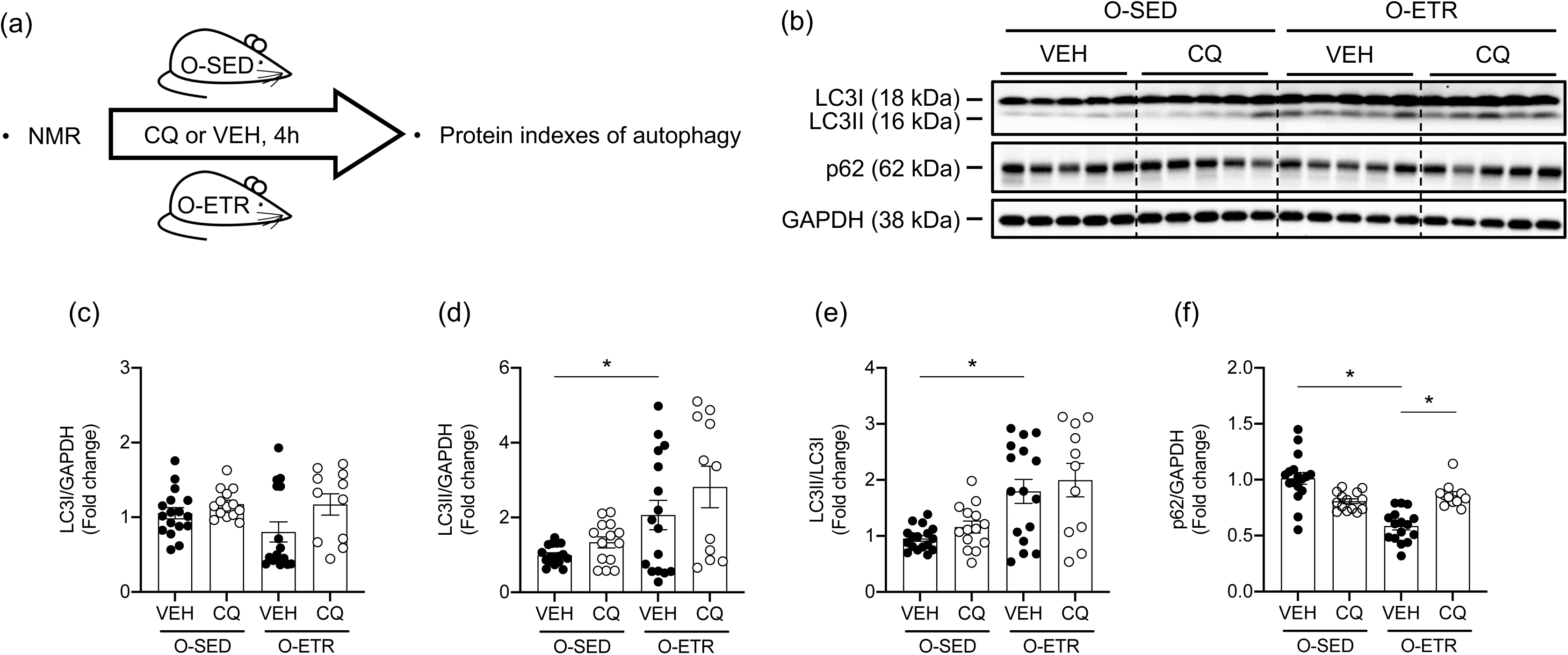
*Autophagic flux is improved in hearts from trained vs. untrained mice.* (a) Body composition was assessed using TD-NMR in older mice (O, 24-mo) that did (ETR) or did not (SED) train for 12-weeks. Vehicle (VEH; phosphate buffered saline) or chloroquine (CQ; 75 mg CQ / g of lean body mass) was administered IP to cohorts of O-SED and O-ETR mice. Four h later animals were anesthetized, hearts were excised, and segments of myocardium were prepared for immunoblotting. Representative images (b) and mean data ± standard error are shown (c-f). LC3-II (d) normalized to GAPDH and LC3-II / LC3I were higher in myocardium from O-ETR vs O-SED. p62 (f) normalized to GAPDH were less in myocardium from O-ETR vs. O-SED mice treated with VEH, whereas LC3-II:GAPDH and LC3-I:LC3-II were similar between groups. CQ treatment increased p62:GAPDH in myocardium from ETR but not SED mice (f), whereas LC3-I, LC3-II, and LC3-II:LC3-I normalized to GAPDH were not affected (c-e). For c-f, n=11- *p<0.05 vs O-SED; #p<0.05 vs O-ETR. Data are expressed as fold change relative to values obtained from O-SED-VEH mice.

### 2.5 Protein aggregate removal, ubiquitinated protein clearance, and lipid peroxidation are improved in the heart by late-in-life exercise training

Bolstering our findings shown in Figure S2, protein aggregates (Figure 5a-c), ubiquitinated proteins (Figure 5d,e), and indexes of lipid peroxidation (Figure 5f,g), were elevated (p < .05) in hearts from older-SED vs. adult-SED mice (histogram 3 vs. 1). Importantly, each age-associated disruption was lessened (p < .05) in older-ETR vs. older-SED mice (histogram 4 vs. 3; Figure 5a-g). Adult mice were refractory to the effects of exercise-training concerning protein aggregates, ubiquitinated proteins, and lipid peroxidation (Figure 5 b,c,e,g; histogram 1 vs. 2). mRNA expression of the antioxidant enzymes SOD1, SOD2, and catalase were elevated by exercise training in aged mice (Figure 5h).

**Figure 5.**
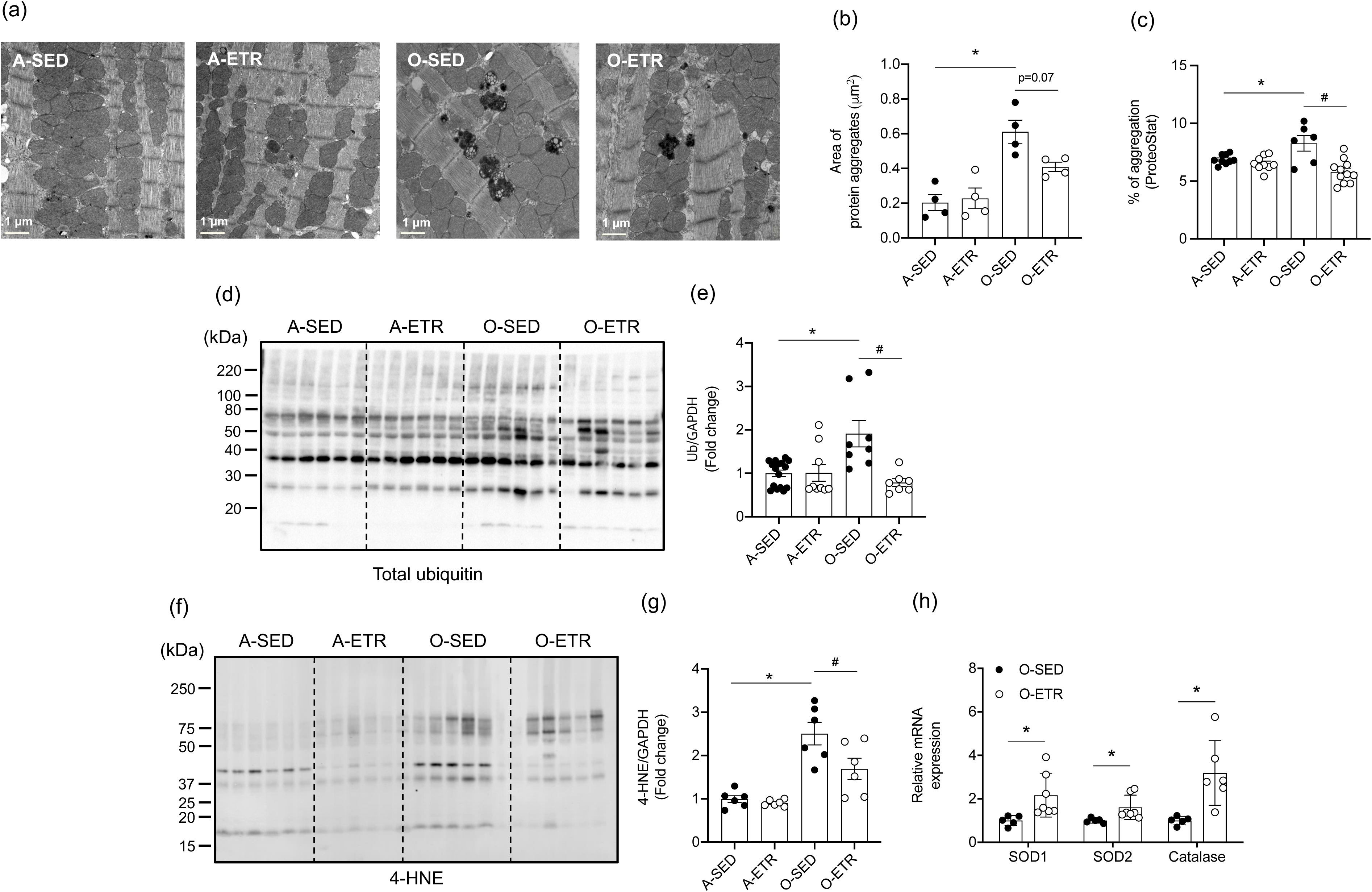
*Protein aggregate accrual, ubiquitinated protein accumulation, and lipid peroxidation observed in hearts from older vs. adult mice is ameliorated by exercise-training.* Adult (A, 5 mo) and older (O, 21 mo) mice did (ETR) or did not (SED) complete 12-weeks of treadmill-running (Figure 3a). At least 24 h following the last exercise bout, A (8 mo) and O (24 mo) mice were anesthetized and segments of myocardium were prepared to quantify protein aggregate accumulation via electron microscopy (EM, a-c), ubiquitinated proteins via immunoblotting (IB, d, e), and lipid peroxidation via IB for 4-HNE (f, g). Representative images (a,d,f) and mean data ± standard error are shown (b,c,e,g). Indexes of protein accrual (a-c), total ubiquitin (d,e), and 4-HNE (f,g) otherwise elevated in hearts from O-SED vs. A-SED mice were less in O-ETR vs. O-SED mice. Superoxide dismutase 1 (SOD1) and 2 (SOD2) and catalase, were measured via qPCR in hearts from A and O mice (h). Representative EM images were photographed at x 2700 magnification. For (b), n=4 mice per group, n=8-16 fields of view; (c), n=6-11; (e), n=7-15; (g), n=6; (h), n=5-7. *p<0.05 vs A-SED; #p<0.05 vs O-SED. For (h), *p<0.05 vs O-SED. Data are expressed as fold change relative to values obtained from A-SED mice.

### 2.6 Exercise training rejuvenates myocardial function in older mice

Next we determined whether late-in-life, training-induced improvements in steady-state autophagy, protein clearance, and oxidant stress associated positively with myocardial function. Indexes of systolic (Figure 6b-e) and overall LV function (i.e., MPI; Figure 6j) improved (p < .05), whereas estimates of diastolic function were unchanged (Figures 6f-h and S8), in old-ETR vs. old-SED mice. A similar pattern of results was observed in adult-ETR vs. adult-SED animals (Figure S9, S10). Importantly, in older mice, the **training**-evoked increase in p62 protein degradation (indicating greater autophagy) associated positively (p <.028) with the training-induced lowering of MPI (indicating greater LV function; Figure 6k). Collagen type 1 area (Figure S11a,b), cardiomyocyte area (Figure S11a,c), and mRNA expression of profibrotic genes (Figure S11d), were similar between old-SED and old-ETR mice.

**Figure 6.**
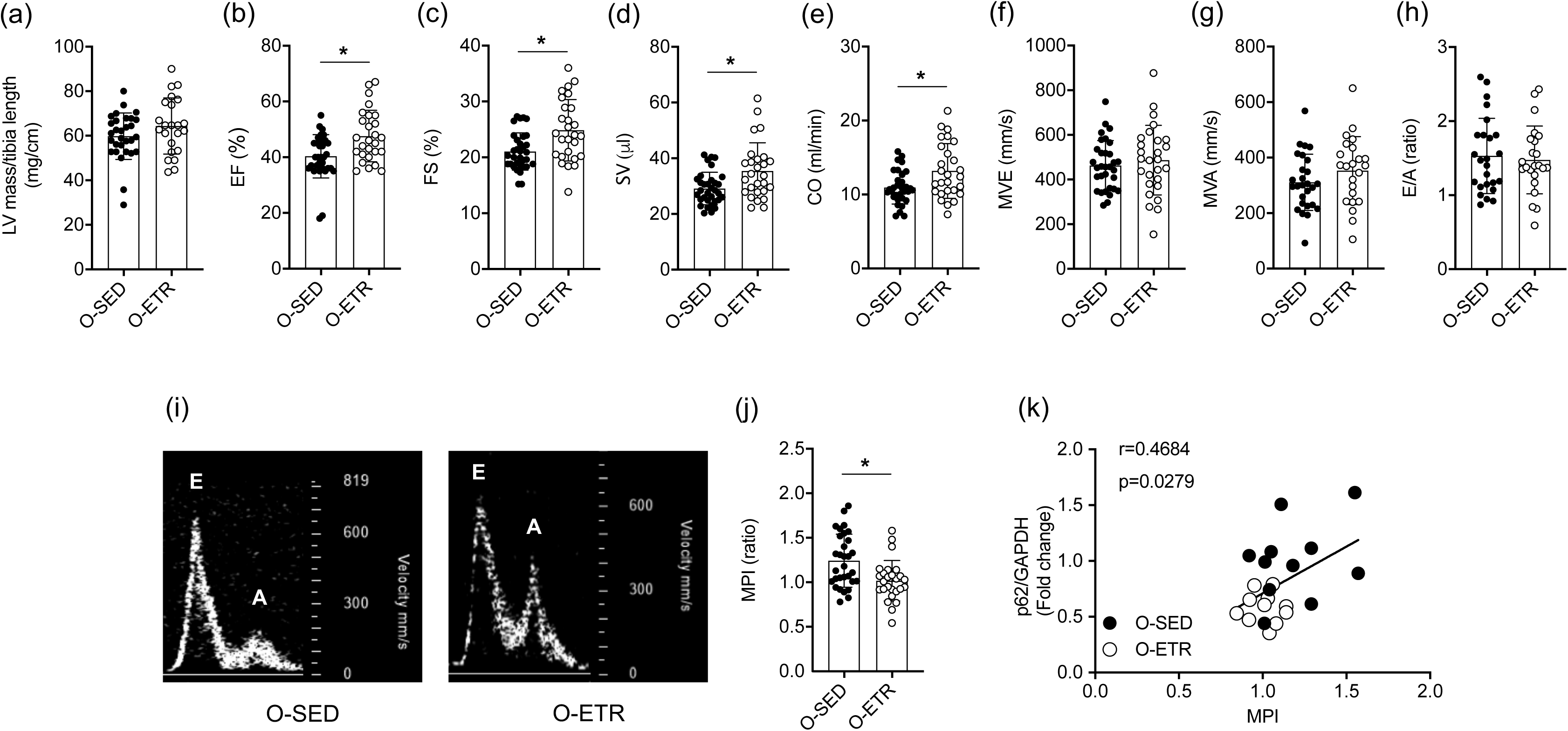
*Exercise-training improves myocardial function in older mice.* Adult (A, 5 mo) and older (O, 21 mo) mice did (ETR) or did not (SED) complete 12-weeks of treadmill-running. At least 24 h following the last exercise bout transthoracic echocardiography was completed on A (8 mo) and O (24 mo) mice. Mean data ± standard error are shown (a-h, j). Left-ventricular mass normalized to tibia length were not different between groups (a). Ejection fraction (EF,%; b), fractional shortening (FS, %; c), stroke volume (SV, μl; d), and cardiac output (μl/min, e) were greater in O-ETR vs. O-SED mice, whereas passive diastolic filling (MVE, mm/s, f), active diastolic filling (MVA, mm/s, g), and the MVA/MVE ratio (E/A, h) were not different between groups. (i) Representative images of blood flow velocity obtained during the assessment of MVE and MVA from both groups. The myocardial performance index (MPI, j), an indicator of overall LV function, was less in O-ETR vs. O-SED mice, indicating that function improved in O-ETR vs. O-SED animals. (k) The correlation between protein expression of p62/GAPDH and MPI was strong in hearts from O-SED and O-ETR mice. For panels (a-h) and (j), n=27-33, *p<0.05 vs O-SED. Data are expressed as mean ± SEM. For (k), n=11.

## 3 DISCUSSION

Our primary aim was to test the hypothesis that late-in-life exercise-training rejuvenates indexes of cardiac autophagy, improves clearance of ubiquitinated proteins, and reestablishes myocardial function. First we substantiated earlier findings that repressed autophagic flux in the myocardium of older vs. adult mice exists, and that this is associated with protein aggregate accrual, oxidant stress, and cardiac dysfunction. Next we demonstrated for the first time that a physiological intervention i.e., progressive resistance treadmill-running, improves autophagic flux, protein clearance, redox balance, and cardiac function, in hearts from older mice. These data indicate positive crosstalk exists between regular physical activity and myocardial autophagy in the context of primary aging.

### Autophagic flux is depressed in hearts from older mice

Repressed autophagy is observed in a wide variety of conditions associated with aging, including neurodegenerative diseases, normal brain aging, osteoarthritis, insulin resistance, atherosclerosis, macular degeneration, suppressed hepatic proteolysis, and endothelial cell dysfunction (Bharath et al., 2017; Marino & Lopez-Otin, 2004; S. K. Park et al., 2019; Rubinsztein et al., 2011; Vittorini et al., 1999). A close examination of the literature reveals that the impact of aging on cardiac autophagy in pre-clinical murine models is not congruent. (Boyle et al., 2011; Hua et al., 2011; Inuzuka et al., 2009; Li et al., 2017; L. Ma et al., 2017; Marin-Aguilar et al., 2020; Peng et al., 2013; Ren et al., 2017; Taneike et al., 2010; Wang et al., 2018; Wu et al., 2016; Zhou et al., 2017; Zhou et al., 2013). On balance, most studies investigating this issue have assessed steady-state autophagy by quantifying protein expression of LC3-II and / or p62. The membrane-bound lipidated form of cytosolic LC3-I i.e., LC3-II accumulates as the phagophore membrane is formed and extended during the process of autophagy. Atg3 is the phosphatidylethanolamine-transferase that performs the final lipid-conjugation modification of LC3 required for completing the conversion of cytosolic LC3-I to membrane-bound LC3-II (Mizushima, 2007). The adaptor protein p62, which tethers targeted cargo destined to become engulfed in the autophagosome, is degraded as autophagy proceeds.

A variety of studies indicate p62 accumulates in hearts from aged vs. adult mice (Li et al., 2020; Ren et al., 2017; Wang et al., 2018; Wu et al., 2016; Y. Zhang et al., 2017), and translational relevance of these findings to older humans was recently reported (Li et al., 2020). Results concerning LC3-II are less clear. With regard to C57BL/6J mice: (i) LC3II / GAPDH (Taneike et al., 2010) and LC3II / LC3I (Hua et al., 2011; Ren et al., 2017) declined in hearts from ∼ 26 mo vs. ∼ 4 mo animals; (ii) LC3II / LC3I increased in 18 mo vs. 2 mo mice; (Boyle et al., 2011) and (iii) LC3II / LC3I was not different between ∼ 23 mo and ∼ 4 mo animals (Li et al., 2020; Wu et al., 2016). We observed increased LC3I:GAPDH, LC3II:GAPDH, and p62:GAPDH in older vs. adult mice from two independent cohorts treated identically (Figure 1, Figure 3). Because Atg3 mRNA and protein expression was lower in both groups of older vs. adult mice (Figure S2, S6), elevated LC3I:GAPDH observed in older animals could have resulted from an inability to perform the lipid conjugation step whereby cytosolic LC3I is converted to membrane-bound LC3II. With regard to LC3II and p62 accrual observed in hearts from older vs. adult mice, this might be secondary to a defect that exists later in the process of autophagy and we tested this (Loos, du, & Hofmeyr, 2014). Separate cohorts of adult and old mice were treated with the lysosomal acidification inhibitor CQ to assess autophagic flux (Glick et al., 2010; Gottlieb et al., 2015; Pires et al., 2017). This approach has been used in the context of cardiac aging on two occasions. Wu et al. treated C57BL/6J mice with the vacuolar H^+^- ATPase inhibitor bafilomycin (0.3 mg/kg IP x 7 days) which impairs lysosomal acidification, blocks autophagosome-lysosome fusion, and thereby prevents degradation of autophagolysosome (Kawai, Uchiyama, Takano, Nakamura, & Ohkuma, 2007; Yoshimori, Yamamoto, Moriyama, Futai, & Tashiro, 1991). Compared to mice that were administered a vehicle-control, bafilomycin increased LC3II : LC3I and p62 protein expression in myocardial lysates from 4 but not 22 mo old mice (Wu et al., 2016). Using a different species and experimental setting, Ma et al. reported that cardiomyocytes isolated from hearts of 4 mo rats displayed greater LC3 puncta and p62 expression after treatment with bafilomycin (100 nM x ∼ 4h) compared to results obtained from ∼24 mo rats (L. Ma et al., 2017). Both studies concluded that constitutive autophagosome formation is robust in myocardium from adult but not older mice and our results are supportive. Specifically, CQ-treatment increased LC3I:GAPDH, LC3II:GAPDH, and p62:GAPDH in hearts from adult but not old mice, underscoring the statement that autophagosome clearance capacity is compromised in aged myocardium (Figure 1).

### Repressed myocardial autophagic flux associates with cardiac proteotoxicity, oxidant stress, and contractile dysfunction

Strong rationale exists that repressed autophagosome formation contributes importantly to accelerated cardiac aging. In a loss of autophagy approach, adult mice with cardiac-selective Atg5 deletion (Taneike et al., 2010), and cardiac-specific mTOR activation via miR-199a overexpression (Li et al., 2017) or tuberous sclerosis complex 1 and 2 depletion (Taneike et al., 2016), exhibit important characteristics of cardiac aging i.e., protein aggregate accrual, interstitial fibrosis, LV hypertrophy, oxidant stress, and / or cardiac dysfunction. In addition to compromised autophagic flux, old vs. adult mice in the present study displayed each one of these features of cardiac aging. Highlighting an association between repressed cardiac autophagy and myocardial dysfunction, elevated cardiac p62 protein expression correlated significantly with a well-accepted estimate of overall LV dysfunction i.e., the MPI (Goroshi & Chand, 2016; Tei et al., 1995) (Figure 2l). Using a gain of autophagy procedure, mice with cardiac-selective Atg7 overexpression (*Atg7* transgenic mice) were crossed with *CryAB^R120G^* mice, a model of desmin-related cardiomyopathy that exhibits impaired autophagic flux together with the accumulation of preamyloid oligomer (PAO), a toxic component in many of the protein misfolding based neurodegenerative diseases (Maloyan, Gulick, Glabe, Kayed, & Robbins, 2007; Pattison, Osinska, & Robbins, 2011). As would be predicted, autophagic flux was greater, and accrual of cytotoxic proteins, impaired cardiac performance, and early mortality was less severe, in *CryAB^R120G^ x Atg7* transgenic mice vs. *CryAB^R120G^* animals (Bhuiyan et al., 2013). Based on previous results using loss of autophagy and gain of autophagy approaches involving the myocardium, it is not unreasonable to suggest that impaired autophagic flux (Figure 1) contributed importantly to the accrual of ubiquitinated proteins and elevated lipid peroxidation (Figure 5, S1), protein aggregate accumulation (Figure 5), increased fibrosis (Figure S4), and systolic and diastolic dysfunction (Figure 2, S3) exhibited by older vs. adult hearts in our study. While precise mechanisms responsible for the age-associated reduction in Atg3 in cardiomyocytes have not been reported, evidence exists that oxidative stress inhibits Atg3 enzyme function in HEK 293 cells (Frudd, Burgoyne, & Burgoyne, 2018) and Atg3 protein expression in mouse brain endothelial cells (Kamat, Kalani, Tyagi, & Tyagi, 2015).

### Late-in-life interventions attenuate cardiac dysfunction by activating autophagy

Although genetic manipulations (e.g., Atg7 overexpression) cannot be used clinically to upregulate autophagy in conditions associated with cardiac proteotoxicity at present, benefits from inducing this protein degradation pathway late-in-life via nutraceutical (e.g., spermidine), lifestyle (e.g., caloric restriction) and pharmacological (e.g. rapamycin) maneuvers have been demonstrated (Eisenberg et al., 2016; Flynn et al., 2013; Sheng et al., 2017). For example, cardiac hypertrophy was attenuated and diastolic function was preserved in C57BL/6J mice that consumed spermidine-supplemented vs. vehicle-treated water from 18-24 months of age. The autophagy-boosting effect of spermidine shown originally in flies and yeast (Eisenberg et al., 2009; Morselli et al., 2011) was confirmed in the myocardium, and cardiac-selective Atg5 deficient animals were refractory to this intervention, substantiating that autophagy is necessary for the benefits of this polyamine to be observed (Eisenberg et al., 2016). Sheng et al. fed one group of C57BL/6J mice standard chow ad libitum, whereas another cohort consumed 40% fewer calories from the same diet, from 19-22 months of age (Sheng et al., 2017). AMPK is phosphorylated and activated by nutrient insufficiency to an extent that increases expression of autophagy-associated genes (Kubli & Gustafsson, 2014; Sanchez et al., 2012) and this was confirmed in myocardium from calorie-restricted vs. ad libitum fed mice. Age-associated cardiac fibrosis, LV hypertrophy, and compromised ejection fraction were less severe in mice that ingested calorie-restricted vs. standard diet from 19-22 months of age i.e., late-in-life. While inhibiting TOR (target of rapamycin) signaling extends lifespan in organisms from worms to mice (Harrison et al., 2009; Kapahi et al., 2004; Vellai et al., 2003), Flynn et al. first reported the cardiovascular benefits of this intervention in the context of aging (Flynn et al., 2013). Mice treated with rapamycin from 24-27 months of age had repressed pro-inflammatory signaling in the myocardium, less LV hypertrophy, and preserved systolic function vs. age-matched controls. Although rapamycin repressed mTORC1 signaling in the myocardium as demonstrated by reduced pS6K : total S6, evidence for upregulated cardiac autophagy *per se* was not presented. While each of these approaches to boost autophagy in the context of aging attenuated indexes of cardiac dysfunction (Eisenberg et al., 2016; Flynn et al., 2013; Sheng et al., 2017), it is unknown if this benefit associated positively with improved autophagic flux and protein clearance in the myocardium because neither of these endpoints was assessed.

A lifestyle intervention with potential to improve autophagy, clear damaged proteins, and beneficially influence the aging-associated decline in cardiac function is dynamic exercise. He et al. first showed in mice that 30-80 -min treadmill-running increases LC3-GFP puncta, LC3II:LC3I, and p62 degradation in the heart (He et al., 2012). Beta cell lymphoma/leukemia 2 (Bcl-2) is an anti-apoptotic and anti-autophagy protein that inhibits autophagy by directly interacting with beclin 1 at the endoplasmic reticulum (He & Levine, 2010). The authors reported that the Bcl-2–beclin-1 complex dissociates in response to treadmill-running, and this finding was confirmed by Bhuiyan et al. in mice that completed an acute bout of voluntary wheel running (VWR) (Bhuiyan et al., 2013). Because long-term VWR decreased the amyloid load in mice with neurodegenerative disorders e.g., Alzheimers disease (Lazarov et al., 2005) the Robbins laboratory group sought to determine whether this form of “environmental enrichment” might initiate myocardial autophagy to an extent that improves cardiac proteostasis. Providing strong proof of concept, the authors reported a 47% reduction in cardiac PAO accumulation in *CryAB^R120G^* mice that completed 6-months of VWR vs. *CryAB^R120G^* animals that did not train, but indexes of autophagy were not assessed (Maloyan et al., 2007). The same investigative team later demonstrated that VWR increased mRNA expression of Atg4, Atg5, and Wipi1 in myocardium from *CryAB^R120G^* mice vs. untrained mice, but neither autophagic flux nor protein aggregate accrual were assessed (Bhuiyan et al., 2013). In the latter study it is extremely interesting to note that functional endpoints assessed via echocardiography (LVIDs, LVIDd, EF) appear identical between *CryAB^R120G^ x Atg7* transgenic mice and *CryAB^R120G^* mice that completed VWR, suggesting that exercise-training conferred benefits similar to genetic autophagy activation and vice versa.

In light of the interesting findings from interventions involving nutraceuticals, pharmaceuticals, and lifestyle alterations (e.g., caloric restriction, regular physical activity) in older mice, we tested whether late-in-life exercise training induces cardiac autophagy to an extent that improves proteostasis and lessens myocardial dysfunction. As anticipated, the intensity, frequency, and duration of “forced” exercise training produced functional and biochemical evidence indicating our protocol was efficacious (Figure S5). In support of our hypothesis, aging-associated increases in LC3I:GAPDH and p62:GAPDH were lowered in hearts from old-ETR vs. old-SED mice (Figure 3c,f; Figure 4c,f). Importantly, CQ-induced p62:GAPDH accumulation occurred in hearts from old-ETR but not old-SED mice (Figure 4f). Collectively, these results indicate that 3-mo treadmill-running improves autophagic flux in hearts from older mice, and these observations are concurrent with reduced myocardial protein aggregates (Figure 5a, b, c), ubiquitinated proteins (Figure 5d, e), and lipid peroxides (i.e., 4- HNE; Figure 5f, g). It is not unreasonable to suggest that elevated Atg3 in hearts from old-ETR vs. old-SED mice (Figure S7b) facilitated the conversion of cytosolic LC3I to LC3II to thus lower LC3I:GAPDH. While the mechanism(s) responsible for elevated mRNA and protein expression of Atg3 in old-ETR vs. old-SED mice is unclear and has not been reported earlier, improved redox balance (Figure 5) displayed by hearts from old-ETR mice might be responsible, based on results from studies using other cell types (Frudd et al., 2018; Kamat et al., 2015). The ability of CQ to promote accrual of p62 in hearts from old-ETR but not old-SED mice (Figure 4f) indicates improved autophagic flux and likely explains the training-induced reduction of p62:GAPDH we observed in two separate cohorts of old-ETR vs. old-SED that were untreated (Figure 3f) or treated with vehicle (Figure 4f). Three mo of treadmill-training did not prevent aging-associated cardiac fibrosis (Figure S11a-d). However, multiple indexes of systolic performance, together with a doppler-derived measure of overall LV function that incorporates the time intervals of mitral valve inflow and aortic valve outflow (i.e., MPI)(Tei, Nishimura, Seward, & Tajik, 1997), improved in trained vs. untrained older mice (Figure 6). Highlighting the association between training-induced benefits concerning cardiac autophagy (i.e., reduced cardiac p62 protein expression) and LV function (i.e., lower MPI), a significant correlation existed between these two endpoints in older trained mice (Figure 6k). On balance, our data indicate that rejuvenated autophagic flux contributes importantly to greater protein clearance and attenuated cardiac dysfunction in the context of aging.

We observed a strong association between the: (i) age-related accrual of p62:GAPDH (repressed autophagy) and the increase (i.e., worsening) of MPI (Figure 2l); and the (ii) training-induced reduction in p62:GAPDH (improved autophagy) and the decrease (i.e., improvement) of MPI (Figure 6k). While these findings indicate training-induced elevations in autophagic flux associate positively with preserved myocardial function in the context of primary aging, evidence exists that an 8-week exercise program preserves autophagic flux in adult rats in the setting of a common age-related pathology e.g., heart failure (Campos et al., 2017). Specifically, 12-weeks following myocardial infarction induced heart failure via left anterior descending coronary artery ligation, indexes of cardiac autophagic flux and myocardial function were improved in rats that completed treadmill-running from weeks 4-12 vs. those that did not train. At present it is unknown whether late-in-life exercise training rejuvenates autophagic flux to an extent that improves tolerance to infarction-induced heart failure, but these studies are ongoing in our laboratory.

## 4 EXPERIMENTAL PROCEDURES

### Animals and housing

Male C57BL/6J mice were obtained from the Jackson Laboratories at 4 months of age, and from the National Institute on Aging rodent colony at 18 months of age. All mice were handled according to Institutional approved procedures documented in protocol number 19-07010.

### Myocardial autophagy

First we tested the hypotheses that static (i.e., basal) myocardial autophagy and / or autophagosome formation (i.e., autophagic flux) are repressed in hearts from 23-24-month vs. 7-8-month old mice. Body composition was assessed using time-domain-nuclear magnetic resonance (TD-NMR; Bruker minispec, Bruker Biospin Corporation)(Bharath et al., 2017; Bharath et al., 2015; Pires et al., 2017). Twenty-four h later, chloroquine (CQ; 75 mg IP/g lean body mass) or vehicle-control (phosphate-buffered saline; PBS)(Gottlieb et al., 2015) was administered to separate cohorts of older and adult mice. Four h later, mice were anesthetized and segments of myocardium were obtained to assess protein indexes of autophagy (Bharath et al., 2017), ubiquitinated proteins, and 4-hydroxy-2-nonenal (4-HNE), an α, β-unsaturated hydroxyalkenal that is an estimate of lipid peroxidation. Protein isolation and immunoblotting analyses were performed as we described (Bharath et al., 2017; Bharath et al., 2015; S.-Y. Park et al., 2016; Pires et al., 2017; Symons et al., 2011; Symons et al., 2009; Q.-J. Zhang et al., 2012; Q.-J. Zhang et al., 2009).

### Myocardial function

It was necessary for us to confirm previous reports that myocardial function observed in adult mice is compromised in older animals (Dai & Rabinovitch, 2009; Dai et al., 2009; Flynn et al., 2013). Separate cohorts of adult and older mice were anesthetized lightly with 1-3% isoflurane anesthesia and transthoracic echocardiography was completed to assess indexes of systolic and diastolic function (Pires et al., 2017; Symons et al., 2011).

### Histology, morphology, mRNA gene expression

Twenty-four – 48 h after assessing myocardial function, mice were anesthetized and hearts were segmented to assess protein indexes of autophagy, histology, morphology, and mRNA expression of pro-fibrotic and antioxidant genes (Bharath et al., 2017; Bharath et al., 2015; S.-Y. Park et al., 2016; Pires et al., 2017; Symons et al., 2011; Symons et al., 2009; Q.-J. Zhang et al., 2012; Q.-J. Zhang et al., 2009).

### Exercise training

Body composition was assessed using TD-NMR in 5-month and 21-month old mice. Twenty-four to 48 h later, mice were familiarized with walking/running on a motorized treadmill (Columbus Instruments). On day 4, a workload capacity evaluation test was completed on each mouse. Total workload was calculated as [body weight (kg) x total running time (min) x final running speed (m/min) x treadmill grade (25%)] (Symons et al., 2011; Symons, Rendig, Stebbins, & Longhurst, 2000). After all mice finished the workload capacity evaluation, they were separated randomly into groups that did not (adult-SED and old-SED) or did (adult-ETR and old-ETR) complete a 3 mo progressive resistance treadmill-running program. After 3 mo, body composition, exercise tolerance (maximal workload capacity), and myocardial function were assessed. Each evaluation was separated by 24 h. Twenty-four h after measuring myocardial function, all mice were anesthetized as described, a blood sample was obtained via cardiac puncture, and the heart was excised and segmented to assess protein indexes of autophagy, histology, morphology, and mRNA expression of pro-fibrotic genes (described earlier), together with protein aggregate accumulation. Soleus muscle also was dissected free from both hindlimbs to assess CS enzyme activity (Sigma-Aldrich)(Symons et al., 2000).

### Cardiac protein aggregation

After determining myocardial protein concentrations (Pierce BCA Protein Assay; ThermoFisher) protein aggregate accrual in the heart was measured using a commercially available kit (Proteostat; Enzo Life Sciences)(Laor et al., 2019; Watanabe et al., 2012). Electron microscopy was used as a second approach to estimate cardiac protein aggregates (DiMemmo et al., 2017; Sung et al., 2015). Details concerning each of these methods is provided in online “Experimental procedures.”

### Statistical analyses

Data are presented as mean ± standard of error of the mean. Significance was accepted when p < .05. To determine normality of the distribution for each data set, GraphPad Prism software was used (Poitras, 2006). An unpaired t-test (e.g., EF in adult vs. old mice) was used, as appropriate, to compare two mean values. Comparison among four means was completed using a one-way ANOVA (e.g., cardiac p62:GAPDH among adult, adult-CQ, old, old-chloroquine). In cases when a significant main effect was obtained, a Tukey post-hoc test was used to determine the location of the differences.

## ACKNOWLEDGMENTS

Support was provided for: JMC by a University of Utah (UU) Graduate Research Fellowship and American Heart Association (AHA) grant 20PRE35110066; KL and CR by an American Physiological Society (APS) Undergraduate Research Fellowship (URF) and the UU Undergraduate Research Opportunities Program (UROP); LT by a UU UROP; RG by UU College of Heath (COH) Seed Grant; SKP by AHA grant 17POST33670663; SB by NIH/NHLBI R01HL149870-01A1; and JDS by AHA16GRNT31050004, NIH RO3AGO52848, and NIH/NHLBI RO1HL141540. Nancy Chandler, MS from the UU Electron Microscopy Core is thanked for her assistance.

## AUTHOR’S CONTRIBUTIONS

JMC, SB, JDS designed the study. JMC, KL, CR, LT, MH, MSLCM, KMP, MF, MB, RG, SKP, JDS performed experiments and/or trained the mice. KW and KC performed echocardiography at the UU Small Animal Ultrasound Facility. JMC, SB, MH, JDS analyzed the data. JMC, SB, and JDS wrote the manuscript.

## DATA AVAILABILITY STATEMENT

The raw data supporting the conclusions of this manuscript will be made available by the authors, without undue reservation, to any qualified researcher.

## SUPPORTING INFORMATION

### EXPERIMENTAL PROCEDURES

#### Animals and housing

Male C57BL/6J mice were obtained from the Jackson Laboratories at 4 months of age and from the National Institute on Aging rodent colony at 18 months of age. Until their use, mice were housed at the AALAC-approved Comparative Medicine Center at the University of Utah under specific guidelines which included a 12-h light and 12-h dark-cycle and temperature-controlled environment (22-23°C). Mice were given standard rodent chow and water ad libitum and animals were checked daily by vivarium staff. Under the guidance and regulation of the Institutional Animal Care and Use Committee at the University of Utah, all mice were handled according to approved procedures documented in protocol number 19-07010. *Myocardial autophagy.* First we tested the hypotheses that static (i.e., basal) myocardial autophagy and / or autophagosome formation (i.e., autophagic flux) are repressed in hearts from 23-24-month vs. 7-8-month old mice. A schematic is shown in Figure 1a. Lean mass, fat mass, and fluid mass were assessed in all mice using time-domain-nuclear magnetic resonance (TD-NMR; Bruker minispec, Bruker Biospin Corporation)(Bharath et al., 2017; Bharath et al., 2015; Pires et al., 2017). Twenty-four h later, the lysosomal acidification inhibitor chloroquine (CQ; 75 mg IP/g lean body mass) or vehicle-control (phosphate-buffered saline; PBS)(Gottlieb et al., 2015) was administered to separate cohorts of older and adult mice. Four h later, mice were anesthetized using 2-5% inhaled isoflurane combined with 100% oxygen. When a stable plane of anesthesia was attained, the chest was opened using aseptic procedures, the heart was exposed, excised, and segmented to assess protein indexes of autophagy (Bharath et al., 2017), ubiquitinated proteins, and 4-hydroxy-2-nonenal (4-HNE), an **α, β-unsaturated hydroxyalkenal that is an estimate of lipid peroxidation.** Protein isolation and immunoblotting analyses were performed as described by us (Bharath et al., 2017; Bharath et al., 2015; S.-Y. Park et al., 2016; Pires et al., 2017; Symons et al., 2011; Symons et al., 2009; Q.-J. Zhang et al., 2012; Q.-J. Zhang et al., 2009). In brief, total protein from each heart was separated by SDS-PAGE (4-20%), transferred onto polyvinylidene difluoride (ThermoFisher), and probed with LC3II, LC3I, p62/SQSTM1, total-ubiquitin, 4-HNE, and GAPDH primary antibodies. Alexa Fluor anti-rabbit 680 (Invitrogen) and anti-mouse 800 (VWR International) served as secondary antibodies. Fluorescence was quantified using the Odyssey imager (LI-COR Biosciences).

#### Myocardial function

It was necessary for us to substantiate previous reports that myocardial function observed in adult mice is compromised in older animals (Dai & Rabinovitch, 2009; Dai et al., 2009; Flynn et al., 2013). A schematic is shown in Figure 2a. Separate cohorts of adult and older mice were anesthetized lightly with 1-3% isoflurane anesthesia combined with 100% oxygen while myocardial function was assessed using transthoracic echocardiography (Pires et al., 2017; Symons et al., 2011). Acquisitions were made using a Vevo 2100 high-resolution unit equipped with a 22-55 MHz transducer (Visual Sonic)(Pires et al., 2017; Symons et al., 2011). An investigator blinded to mouse age performed the analyses and reduced the data using the Vevo-Strain software / Vevo 2100 imaging system. Parasternal long axis images acquired in B-mode and parasternal short axis images at the level of the papillary muscles acquired in M-mode were captured using an MS 550D transducer. Indices of systolic function [stroke volume (SV), ejection fraction (EF), fractional shortening (FS), cardiac output (CO)], together with end-diastolic volume, end-systolic volume, and end-systolic left ventricular (LV) mass were assessed. Parasternal short-axis measures in M-mode were used to estimate LV anterior wall thickness in diastole and systole, LV internal diameter in diastole and systole, and end-diastolic and end-systolic volumes. Using pulse wave doppler, diastolic function was estimated by measuring passive (i.e., early due to pressure gradient; E) and active (i.e, later due to atrial contraction; A) velocity of blood flow through the mitral valve (MV). These results were used to calculate the E/A ratio. The myocardial performance index (MPI), an indicator of global left ventricular function, was calculated after measuring isovolumetric contraction time (ICT), isovolumetric relaxation time (IRT), and ejection time (ET), as (ICT + IRT) / ET (Goroshi & Chand, 2016; Tei et al., 1995). VevoLab analysis software 3.1.0 was used to quantify all measures.

#### Histology, morphology, mRNA gene expression

Twenty-four – 48 h after assessing myocardial function, mice were anesthetized as described, and excised hearts were segmented to assess protein indexes of autophagy (described earlier), histology, morphology, and mRNA expression of pro-fibrotic and antioxidant genes. Segments of frozen tissue were cut and embedded in optimal cutting temperature (OCT) compound (Fisher Scientific). Five sections (5 µm thick) per sample were placed on glass slides, and stored at -80°C. To assess type 1 collagen, sections were air dried for 60-min, fixed in ice cold acetone (Fisher Scientific) for 10 min, blocked for 60-min in 3% BSA blocking buffer, and stained with anti-Collagen I antibody (1:200, Abcam) at 4°C overnight. Slides were washed three times with 1X PBS, and incubated with Alexa Fluor Plus 647 goat anti-Rabbit IgG secondary antibody (1:500, Invitrogen) for 60-min at room temperature. Negative control sections were treated with secondary antibody only. Next, sections were mounted with ProLong™ Gold Antifade Mountant with DAPI (Invitrogen). Ten 20X images were randomly acquired per section with an XM10 Olympus fluorescence camera (Dai & Rabinovitch, 2009; Tei et al., 1995). Image quantification was performed using the CellSens Dimension software (Olympus). Percentage of the collagen area in total heart tissue was quantified by ImageJ software (NIH). To assess cardiac myocyte area, sections were stained with wheat germ agglutinin (WGA, Alexa 488, Thermo Fisher) for 60-min at room temperature, followed by DAPI (Alexa 405) for 5-min at room temperature. Ten 20X images were randomly acquired per section with an XM10 Olympus fluorescence camera, and cardiomyocyte cross sectional area (µm^2^) was quantified using CellSens Dimension software (Olympus)(Pires et al., 2017). To assess mRNA expression of fibrosis, total RNA was extracted from segments of myocardium from adult and old mice using the RNeasy Mini Kit (Qiagen). (Bharath et al., 2017; Bharath et al., 2015; S.-Y. Park et al., 2016; Pires et al., 2017; Symons et al., 2011; Symons et al., 2009; Q.-J. Zhang et al., 2012; Q.-J. Zhang et al., 2009). Total RNA was reverse transcribed to cDNA using the QuantiTect Reverse Transcription Kit (Qiagen). The quantitative gene expression assay was performed with specific primers for fibrillin (Fbn) 1, Fbn2, transforming growth factor-β (Tgfb)1, Tgfb2, connective tissue growth factor (Ctgf), superoxide dismutase (SOD) 1 and SOD2, and catalase. Relative gene expression was normalized to 18S. Primer sequences are shown in Table S2.

#### Exercise training

Our main focus was to determine whether late-in-life exercise-training improves basal cardiac autophagy and trafficking of the autophagosome to the lysosome to an extent that rejuvenates protein aggregate clearance and recovers myocardial function. A schematic is shown in Figure 3a. Body composition was assessed using TD-NMR in 5-month and 21-month old mice. Twenty-four to 48 h later, mice were familiarized with walking/running on a motorized treadmill (Columbus Instruments) for 3 consecutive days x 5-10 min/day. On day 4, a workload capacity evaluation test was completed on each mouse i.e., 1 min x 5 m/min x 25% grade, followed by 1 m/min increases in speed each min until maximal exercise capacity was achieved. Electrical shocks were not used. Mice were encouraged to run by tapping their rear using test tube cleaning brushes. Total workload was calculated as [body weight (kg) x total running time (min) x final running speed (m/min) x treadmill grade (25%)] (Symons et al., 2011; Symons et al., 2000).

After all mice finished the workload capacity evaluation, they were separated randomly into groups that did not (adult-SED and old-SED) or did (adult-ETR and old-ETR) complete a 3 mo progressive resistance treadmill-running program. Adult-SED and old-SED mice ran on the treadmill 1 day/week x 5 m/min x 10% grade for 5-min to maintain familiarization. This was required so that SED mice would be able to complete a second workload capacity evaluation test after 3 mo. Adult-ETR and old-ETR mice trained 6 days/week x 3 mo. The initial intensity corresponded to 70% of their workload capacity e.g., 30 min x 11.4 m/min x 5% grade. Over the next 3 mo, running duration, treadmill speed, and/or treadmill grade were increased every four days to 60 min x 17.4 m/min x 15% grade. After 3 mo, body composition, exercise tolerance (maximal workload capacity), and myocardial function were assessed. Each evaluation was separated by 24 h. Twenty-four h after measuring myocardial function, all mice were anesthetized as described, a blood sample was obtained via cardiac puncture, and the heart was excised and segmented to assess protein indexes of autophagy, histology, morphology, and mRNA expression of pro-fibrotic genes (described earlier), together with protein aggregate accumulation. Soleus muscle also was dissected free from both hindlimbs to assess CS enzyme activity (Sigma-Aldrich)(Symons et al., 2000).

#### Cardiac protein aggregation

After determining myocardial protein concentrations (Pierce BCA Protein Assay; ThermoFisher) protein aggregate accrual in the heart was measured using a commercially available kit (Proteostat; Enzo Life Sciences)(Laor et al., 2019; Watanabe et al., 2012). In brief, samples (3 mg/ml) were added to a 96-well plate together with 2 μl of detection buffer, and incubated for 5-min in the dark. Fluorescence output was measured at an excitation setting of 550 nm and emission filter of 600 nm (Varioskan Lux, ThermoFisher). Protein aggregation (%) was calculated based on an 8 point standard curve ranging from 0-12% aggregated IgG. Electron microscopy was used as a second approach to estimate cardiac protein aggregates (DiMemmo et al., 2017; Sung et al., 2015). Cardiac segments placed in 2.5 % glutaraldehyde at the time of collection were embedded in plastic, sectioned at 0.5 um with glass knives, and further trimmed with a diamond knife. Sections were placed onto 200 mesh copper grids, and subsequently stained with saturated uranyl acetate, which extends preservation, improves contrast of the extracellular matrix, membranes, cytoplasm, and DNA, and facilitates the quantification of protein aggregation (Erickson, Anderson, & Fisher, 1987). Three fields of view were chosen based on the quality of the images. Images were photographed at 1100x, 2700x, 4400x, and 11000x magnification (FEI Tecnai T-12, ThermoFisher). Results were calculated by quantifying the number of protein aggregate clusters at 2700x magnification due to the clear and detailed images obtained using NIH Image J software.

## Supplementary Figure legends

**Supplementary Figure 1.** *Ubiquitinated proteins and lipid peroxides are elevated in hearts from older vs. adult mice.* These data are supplemental to Figure 1. Body composition was assessed using TD-NMR in adult (A; 8-mo) and older (O; 24-mo) mice. Vehicle (VEH; phosphate buffered saline) or chloroquine (CQ; 75 mg CQ / g of lean body mass) was administered IP to A and O mice. Four h later mice were anesthetized, hearts were excised, and segments of myocardium were prepared for immunoblotting. Representative images and mean data ± standard error are shown for poly-ubiquitin (total ubiquitin; a, b) and 4-hydroxy-2-nonenal (4-HNE, c,d) normalized to GAPDH. Total ubiquitin and 4-HNE were elevated in hearts from O vs. A mice, but CQ was without effect in both groups. mRNA expression of SOD1, SOD2 and catalase, was assessed via qPCR in hearts from O vs. A mice (e). For (b) and (d), n=10-21, for (e), n=4-7. *p<0.05 vs A. Data are expressed as fold change relative to values obtained from A mice.

**Supplementary Figure 2.** *Atg3 protein expression is lower in myocardium from O vs. A mice.* These data are supplemental to Figure 1. Body composition was assessed using TD-NMR in adult (A; 8-mo) and older (O; 24-mo) mice. Vehicle (VEH; phosphate buffered saline) or chloroquine (CQ; 75 mg CQ / g of lean body mass) was administered IP to A and O mice. Four h later mice were anesthetized, hearts were excised, and segments of myocardium were prepared for immunoblotting. (a,c) Representative images and (b,d) mean data ± standard error are shown for protein expression of Atg3, Atg5, Atg7 normalized to GAPDH. Atg3 protein expression was lower in hearts from O vs. A mice, whereas Atg7 and Atg5 were similar between groups. Atg3, Atg5, and Atg7 were refractory to CQ treatment in A and O mice. For (b-d), n=9-13, *p<0.05 vs A. Data are expressed as fold change relative to values obtained from A mice.

**Supplementary Figure 3.** *Cardiac function is impaired in older vs. adult mice.* These data are supplemental to Figure 2. Transthoracic echocardiography was performed on adult (A; 8-mo) and older (O; 24-mo) mice. Mean data ± standard error are shown (a-d). (a) Left ventricular (LV) internal dimension in systole (LVIDs, mm), (c) end-systolic volume (ESV, µl), and (d) end-diastolic volume (EDV, µl) were higher in O vs. A mice, whereas (b) LV internal dimension in diastole (LVIDd, mm) was not different between groups. For (a-d) n=12-13, *p<0.05 vs A.

**Supplementary Figure 4.** *Indexes of fibrosis exist in hearts from older vs. adult mice.* These data are supplemental to Figure 2. Twenty-four h after completing transthoracic echocardiography in adult (A; 8-mo) and older (O; 24-mo) mice, animals were anesthetized, hearts were excised, and segments of myocardium were prepared using procedures detailed in the text. (a) Representative immunohistochemistry images and (b,c) mean data ± standard error indicate type 1 collagen (%, b) was greater in hearts from O vs. A mice, whereas cardiomyocyte cross-sectional area (μm^2^, c) was similar between groups. The scale bar represents 50 μm (top) and 20 μm (bottom). (d) mRNA expression of fibrosis-related genes assessed via qPCR indicate fibrillin (Fbn) 1, transforming growth factor-β (Tgfb) 2, and connective tissue growth factor (Ctgf) normalized by 18S, were elevated in hearts from O vs. A mice, whereas Fbn2 and Tgfb1 were similar between groups. For (d) data are expressed as fold change relative to values of A. For (b, c), n=5 mice per group, n = 5 fields of view. For (d), n=4-6 per group. *p<0.05 vs A.

**Supplementary Figure 5.** *Exercise-training is efficacious in adult and older mice.* These data are supplemental to Figure 3. Adult (A, 5 mo) and older (O, 21 mo) mice did (ETR) or did not (SED) complete 12-weeks of treadmill-running. Data shown in a-f are from A and O mice at 8 mo and 24 mo, respectively. O-SED mice displayed (a) greater body mass and (b) fat mass, together with (e) reduced exercise capacity and (f) soleus muscle CS enzyme activity, vs. results from A-SED mice, while (c) lean mass and (d) fluid mass were similar between groups. Exercise training reduced (b) fat mass, and increased (e) exercise capacity and (f) CS activity in A and O mice. Data in a-f are mean ± standard error. For (a-d), n=10-24; (e), n=9-18; (f), n=5-19. *p<0.05 vs A-SED; #p<0.05 vs O-SED.

**Supplementary Figure 6.** *Atg3 protein expression is increased by exercise training in older mice.* These data are supplemental to Figure 3. Adult (A, 5 mo) and older (O, 21 mo) mice did (ETR) or did not (SED) complete 12-weeks of treadmill-running. At least 24 h following the last exercise bout, A (8 mo) and O (24 mo) mice were anesthetized, hearts were excised, and segments of myocardium were prepared for immunoblotting. Representative images (a) and mean data ± standard error are shown (b-d). (b) Protein expression of Atg3 normalized to GAPDH was lower in O-SED vs. A-SED mice, whereas (c) Atg5 and (d) Atg7 were similar between the two SED groups. These data recapitulate findings from A and O mice shown in Supplementary Figure 2. (b) Protein expression of Atg3 normalized to GAPDH increased in O- ETR vs. O-SED mice, whereas Atg5 and Atg7 were refractory to exercise-training. Atg3, Atg5, and Atg7 were similar between A-SED and A-ETR animals. Results in panels b-d are expressed as fold change relative to values obtained from A-SED mice. For panels (b-d), n=9-10, *p<0.05 vs A-SED; #p<0.05 vs O-SED.

**Supplementary Figure 7.** *Autophagic flux is improved in hearts from trained vs. untrained older mice.* These data are supplemental to Figure 4. Body composition was assessed using TD-NMR in older mice (O, 24-mo) that did (ETR) or did not (SED) train for 12-weeks. Vehicle (VEH; phosphate buffered saline) or chloroquine (CQ; 75 mg CQ / g of lean body mass) was administered IP to cohorts of O-SED and O-ETR mice. Four h later animals were anesthetized, hearts were excised, and segments of myocardium were prepared for immunoblotting. Representative images (a) and mean data ± standard error are shown (b-d). Atg3, but not Atg5 or Atg7 protein expression normalized to GAPDH, increased in O-ETR vs. O-SED mice. These findings recapitulate those shown in supplemental Figure 6. CQ administration increased Atg3 in O-ETR but not O-SED mice. Atg5 and Atg7 were not responsive to CQ treatment in either O-ETR or O-SED mice. For panels (b-d), n=7-10, *p<0.05 vs O-SED; #p<0.05 vs O-ETR. Data are expressed as fold change relative to values obtained from O-SED mice. Recap data in supp fig 6.

**Supplementary Figure 8.** *Cardiac function in trained and untrained older mice.* These data are supplemental to Figure 3. Adult (A, 5 mo) and older (O, 21 mo) mice did (ETR) or did not (SED) complete 12-weeks of treadmill-running. At least 24 h following the last exercise bout, transthoracic echocardiography was completed on A (8 mo) and O (24 mo) mice. Mean data ± standard error are shown (a-d). (a) Left ventricular (LV) internal dimension in systole (LVIDs, mm) and (b) diastole (LVIDd, mm), and (c) end-systolic (ESV, µl) and (d) end-diastolic (EDV, µl) volume were similar in O-SED and O-ETR mice. For (a-d), n=27-33, *p<0.05 vs O-SED.

**Supplementary Figure 9.** *Exercise-training improves systolic myocardial function in adult mice.* These data are supplemental to Figure 3. Adult (A, 5 mo) mice did (ETR) or did not (SED) complete 12-weeks of treadmill-running. At least 24 h following the last exercise bout transthoracic echocardiography was completed on A (8 mo) mice. Mean data ± standard error are shown (a-h and j). Left-ventricular mass normalized to tibia length was similar between A-SED and A-ETR (a). Ejection fraction (EF,%; b), fractional shortening (FS, %; c), stroke volume (SV, μl; d), and cardiac output (CO, μl/min, e) were greater in A-ETR vs. A-SED mice, whereas passive diastolic filling (MVE, mm/s, f), active diastolic filling (MVA, mm/s, g), and the MVA/MVE ratio (E/A, h) were not different between groups. (i) Representative images of blood flow velocity obtained during the assessment of MVE and MVA from both groups. The myocardial performance index (MPI, j), an indicator of overall LV function, was similar between A-ETR vs A-SED. For (a-h and j), n=8-9, *p<0.05 vs A-SED. Data are expressed as mean ± SEM.

**Supplementary Figure 10.** *Cardiac function in trained and untrained adult mice.* These data are supplemental to Figure 3. Adult (A, 5 mo) mice did (ETR) or did not (SED) complete 12-weeks of treadmill-running. At least 24 h following the last exercise bout, transthoracic echocardiography was completed on A (8 mo) mice. Mean data ± standard error are shown (a-d). (a) Left ventricular (LV) internal dimension in systole (LVIDs, mm) and (b) diastole (LVIDd, mm), and (c) end-systolic (ESV, µl) and (d) end-diastolic (EDV, µl) volume were similar in A-SED and A-ETR mice. For (a-d), n=8-9,

**Supplementary Figure 11.** *Indexes of fibrosis are similar in hearts from trained and untrained older mice.* These data are supplemental to Figure 3. Twenty-four h after completing transthoracic echocardiography in trained (ETR) and untrained (SED) older (24-mo) mice, animals were anesthetized, hearts were excised, and segments of myocardium were prepared using procedures detailed in the text. (a) Representative immunohistochemistry images and (b,c) mean data ± standard error indicate type 1 collagen (%, b) and cardiomyocyte cross-sectional area (μm^2^, c) were similar between groups. The scale bar represents 50 μm (top) and 20 μm (bottom). (d) mRNA expression of fibrosis-related genes assessed via qPCR indicate no differences exist between groups concerning the profibrotic genes fibrillin (Fbn) 1 and 2, transforming growth factor-β (Tgfb) 1 and 2, and connective tissue growth factor (Ctgf), normalized by 18S. For panel (d), data are expressed as fold change relative to values of O-SED. For (b, c), n=6 mice per group, n=5 fields of view. For (d), n=5-9 per group.

