## Supplemental figures and tables for "Late-in-life treadmill-training rejuvenates autophagy, protein aggregate clearance, and function in mouse hearts"

Cho et al, Supplemental Figure 1.

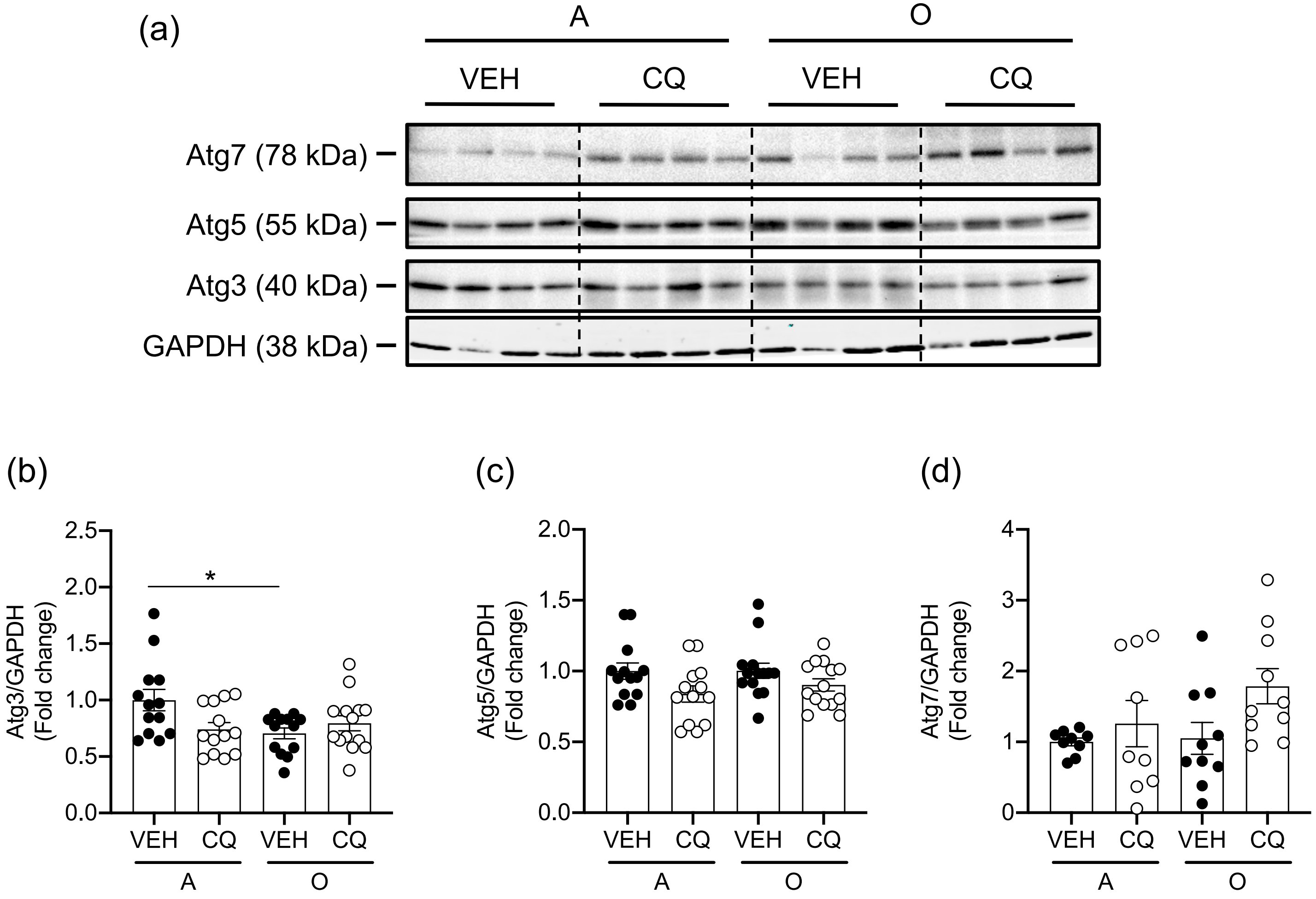

Cho et al, Supplemental Figure 2.

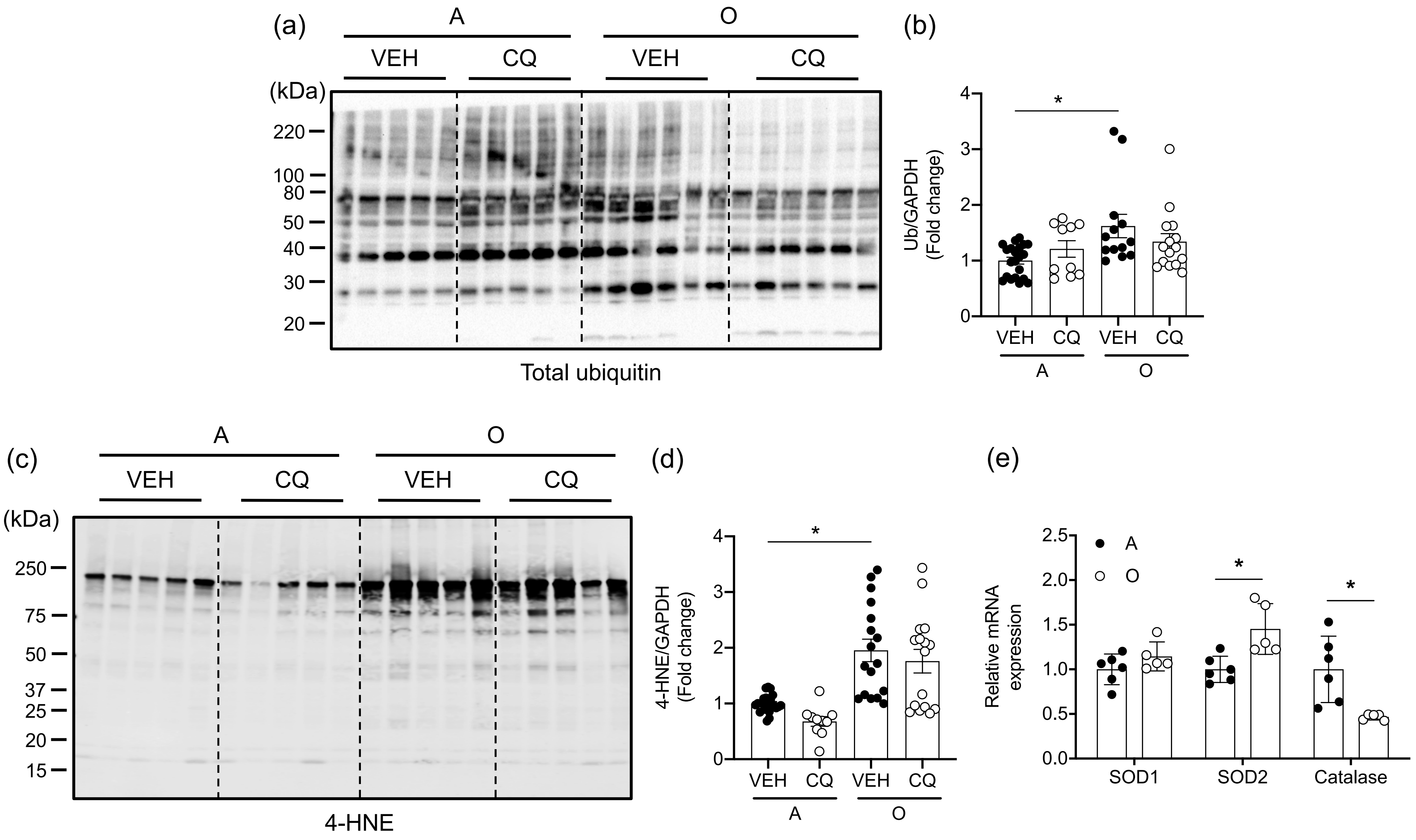

Cho et al, Supplemental Figure 3.

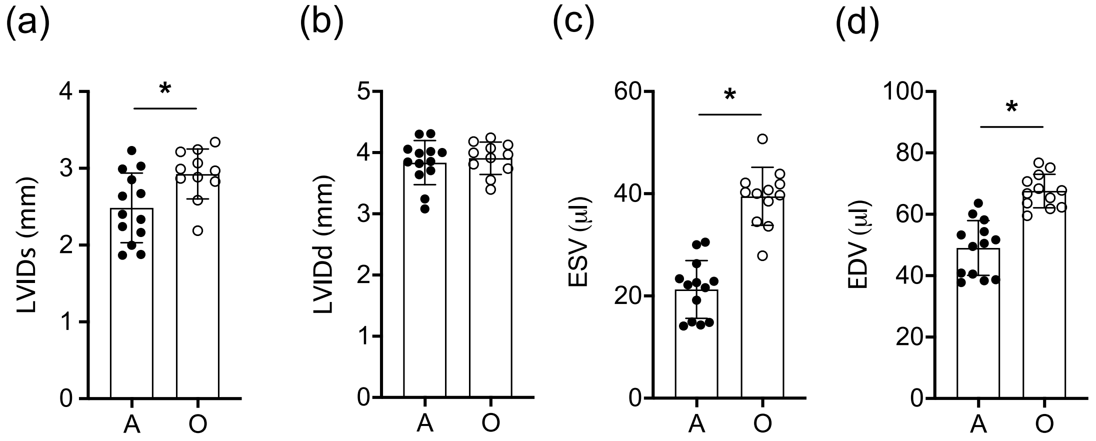

Cho et al, Supplemental Figure 4.

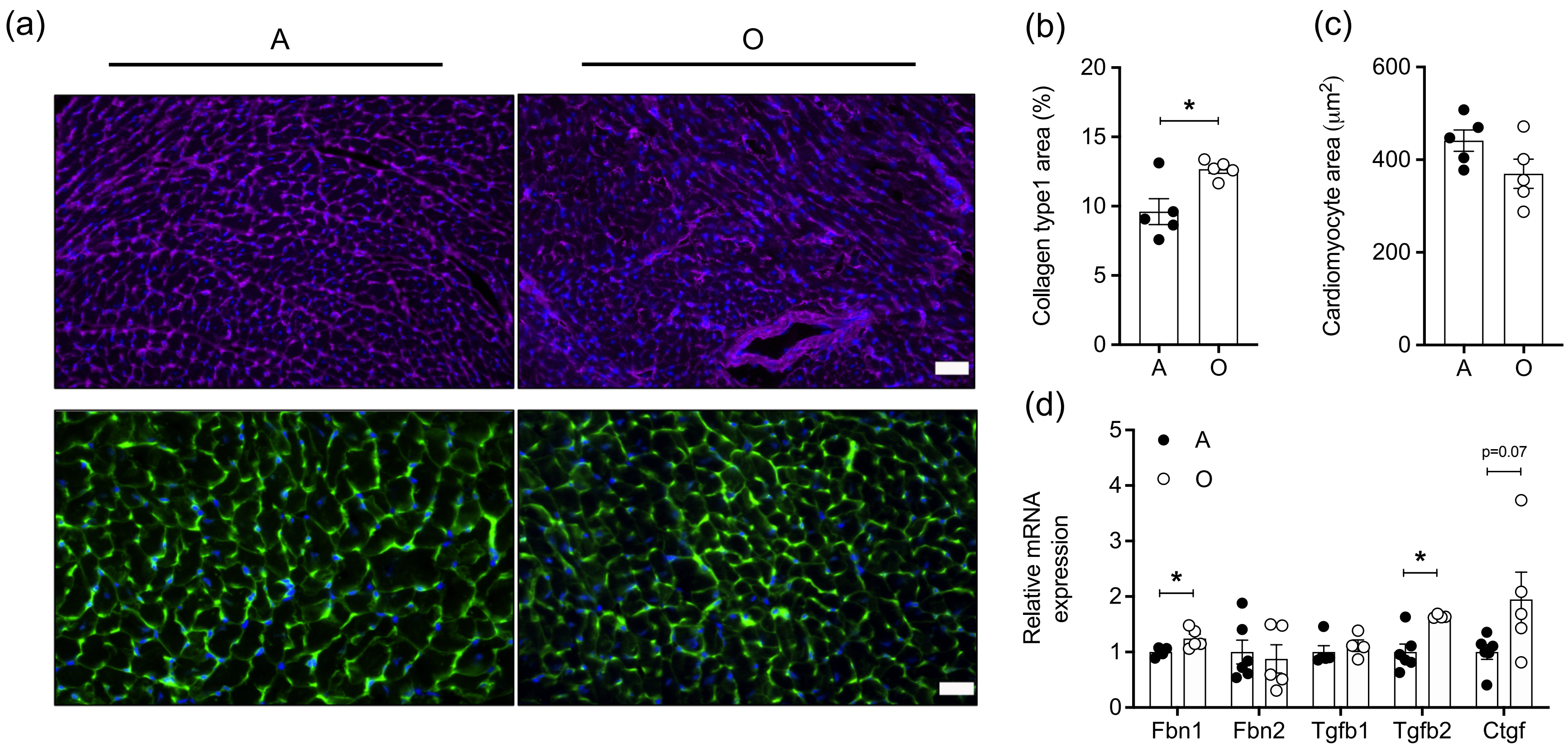

Cho et al, Supplemental Figure 5.

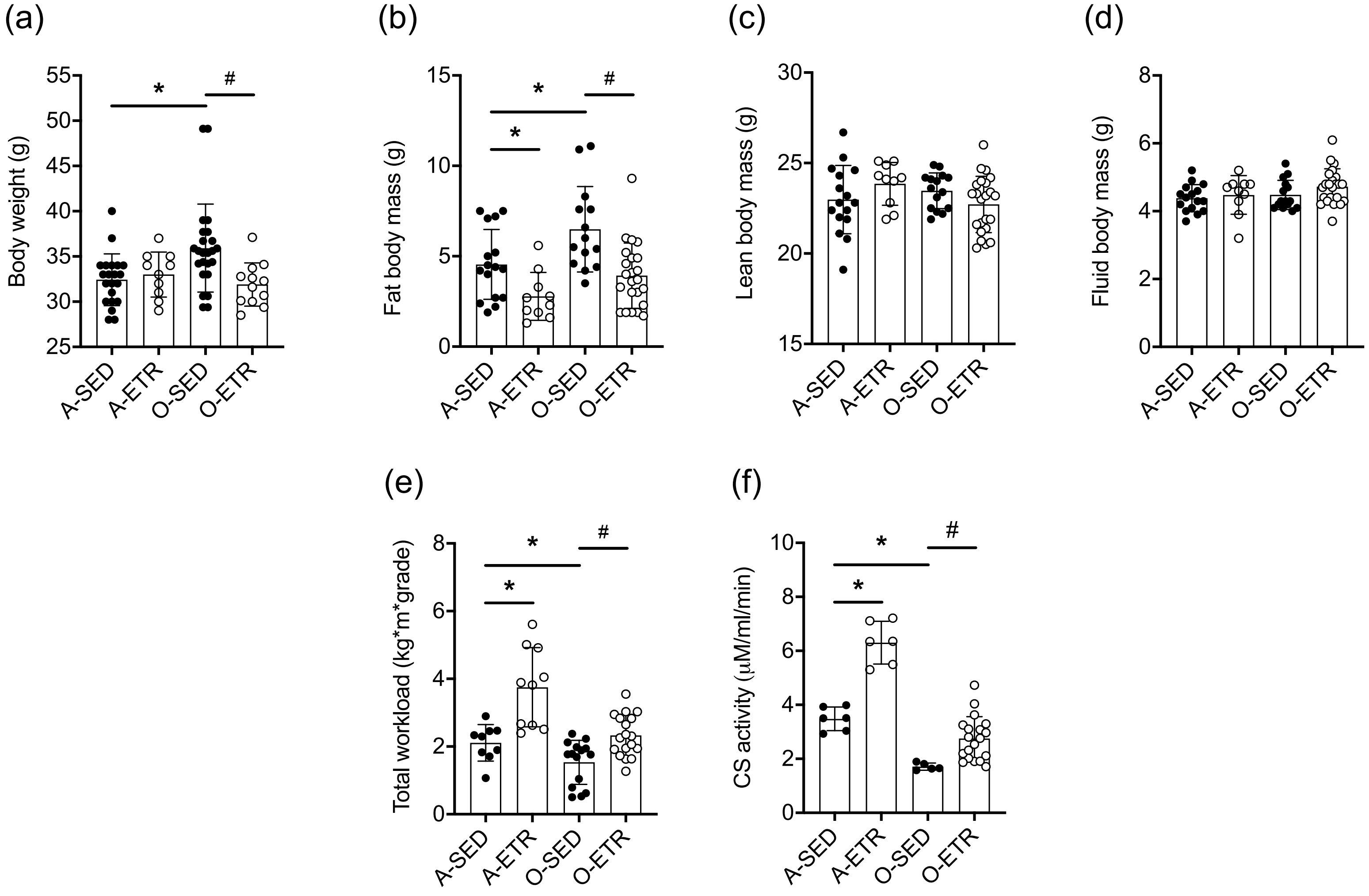

Cho et al, Supplemental Figure 6.

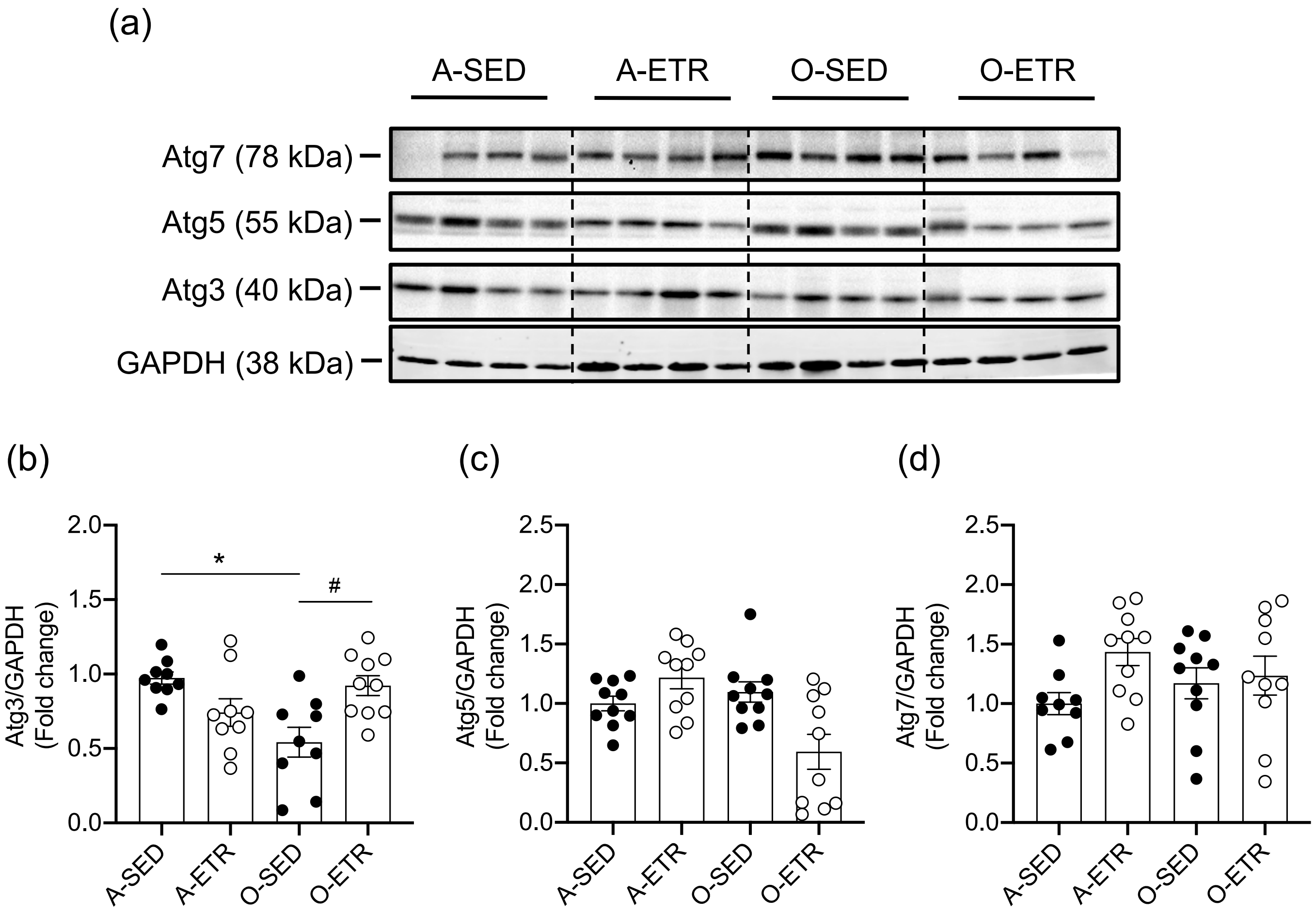

Cho et al, Supplemental Figure 7.

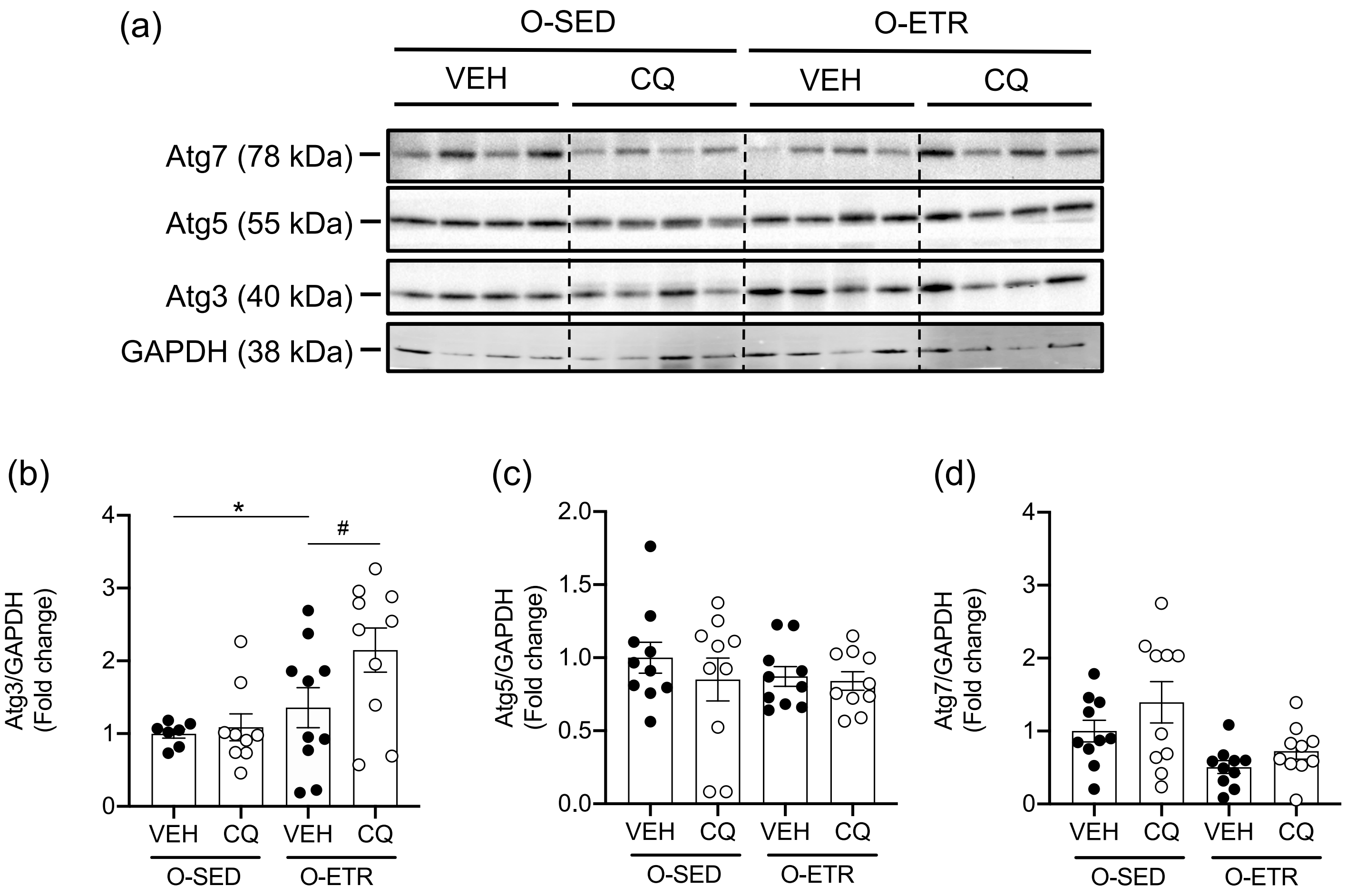

Cho et al, Supplemental Figure 8.

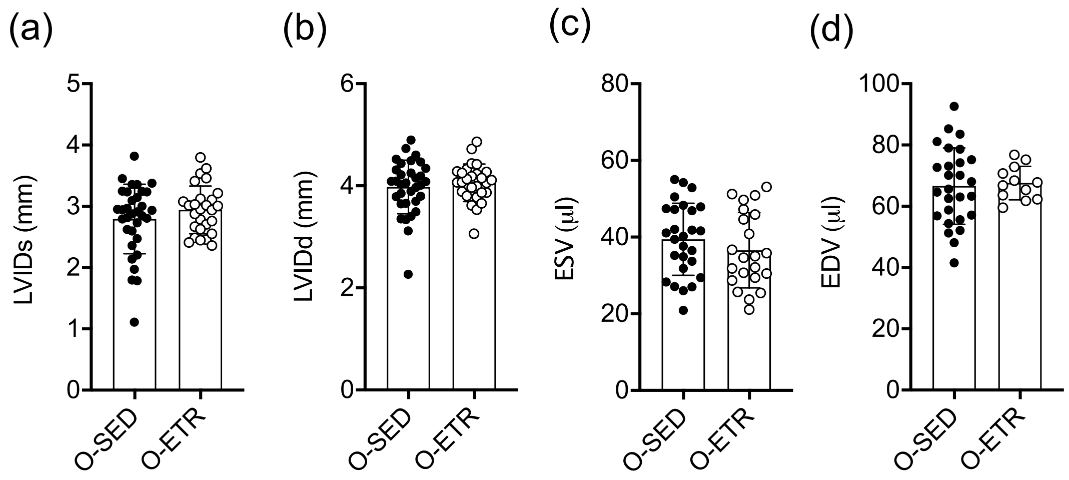

Cho et al, Supplemental Figure 9.

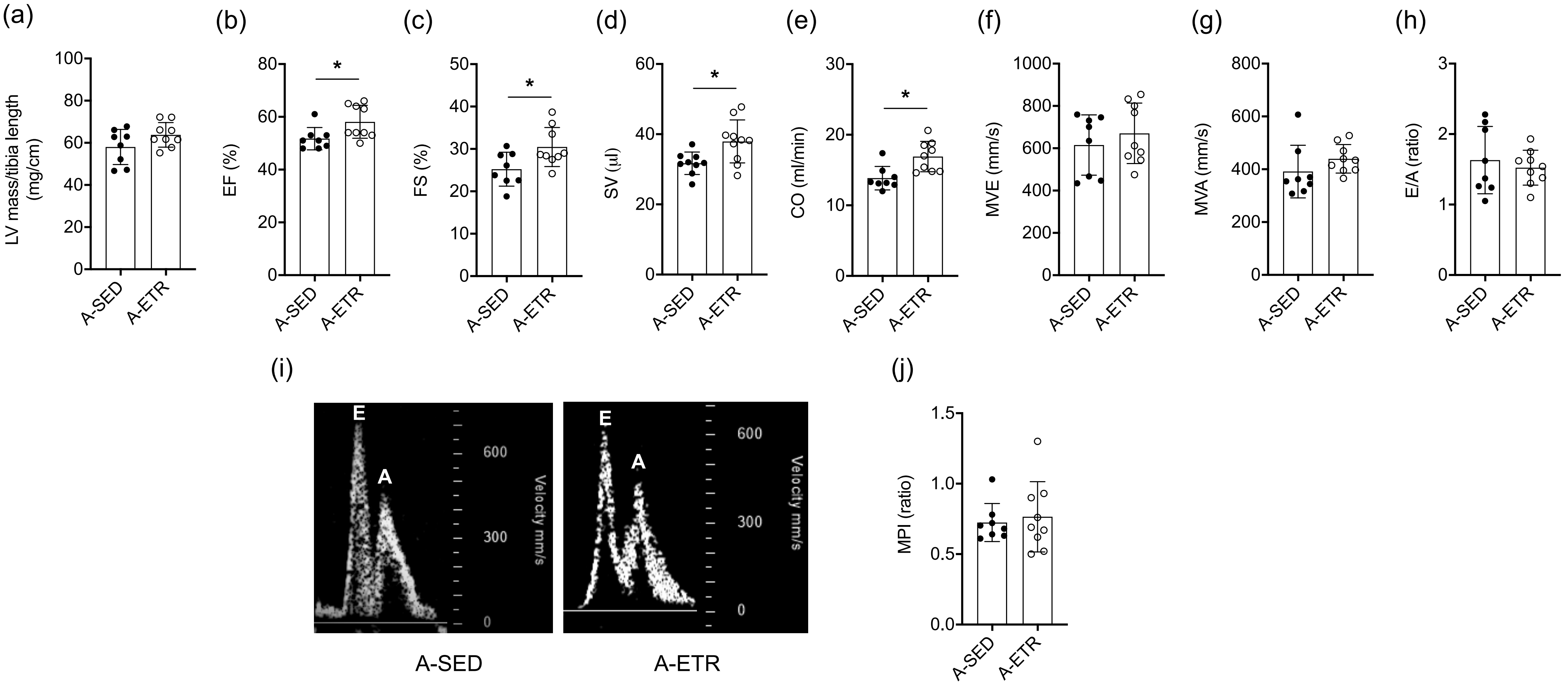

Cho et al, Supplemental Figure 10.

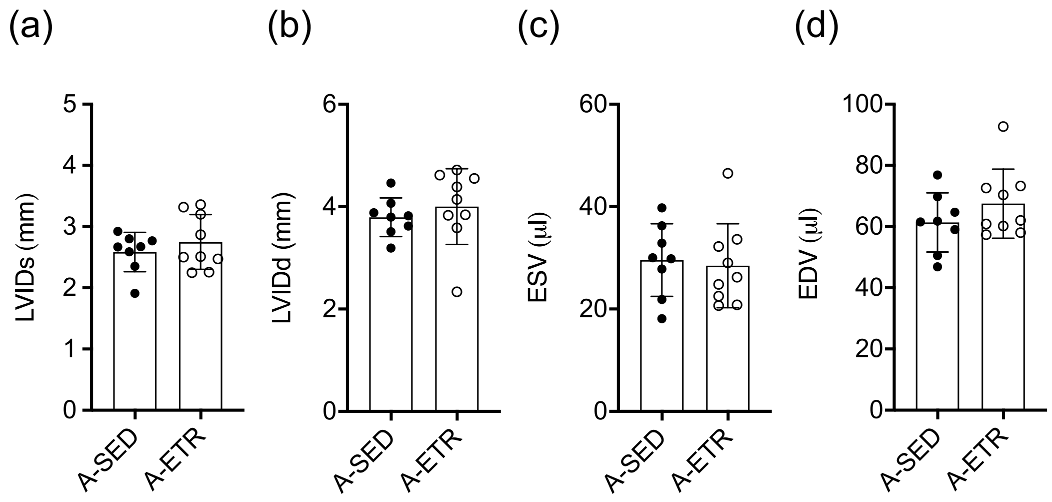

Cho et al, Supplemental Figure 11.

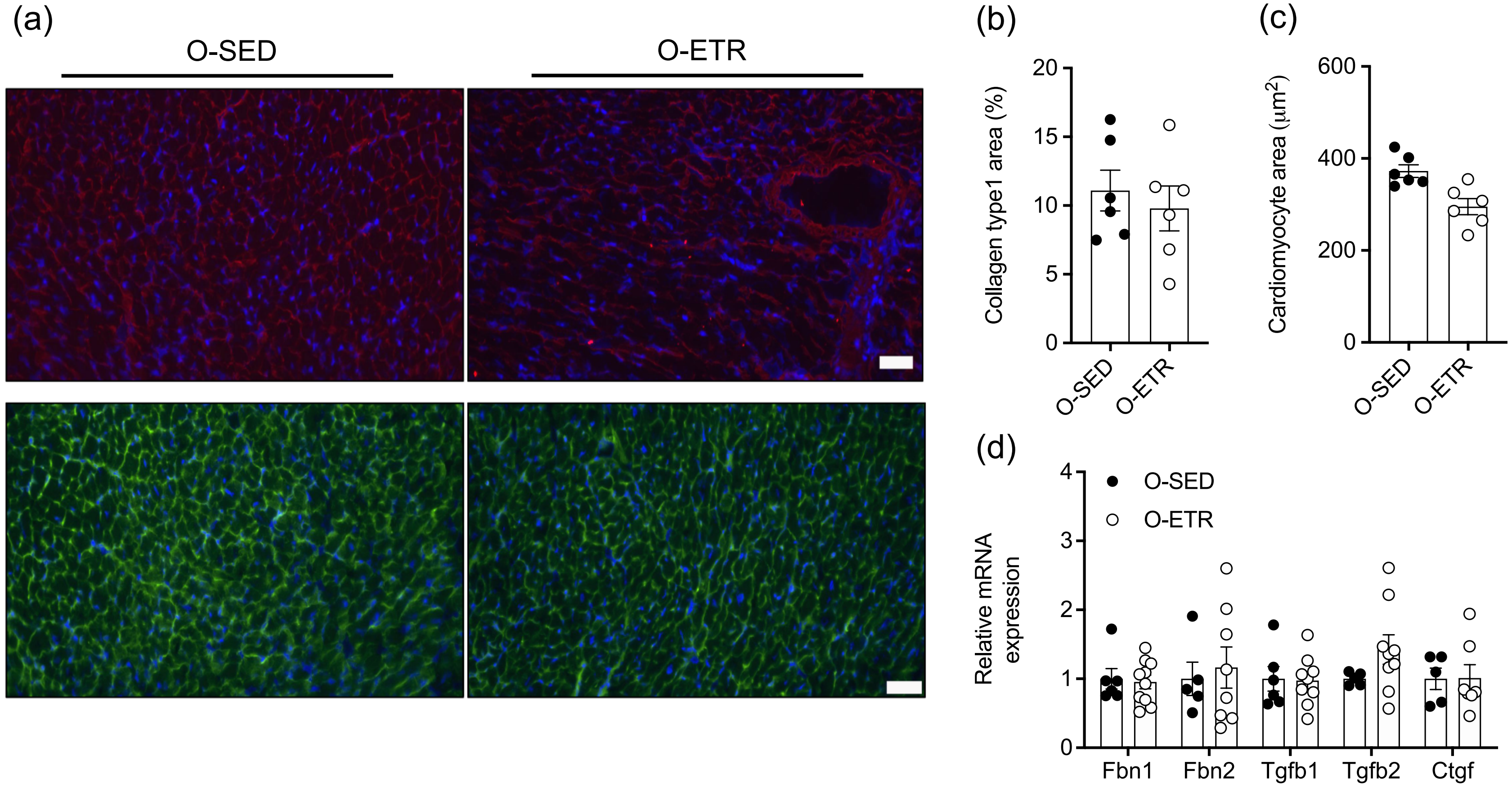

Supplementary Table 1. Animal characteristics – need tables to be completed with table legends+

| | A-SED | A-ETR | O-SED | O-ETR |
|---|---|---|---|---|
| n | 30 | 20 | 40 | 24 |
| Days old at time of study | 228 ± 5 | 222 ± 8 | 677 ± 4 | 728 ± 6 |
| Days of training | - | 78 ± 0 | - | 77 ± 1 |
| Heart weight, g | 0.166 ± 0.004 | 0.186 ± 0.010* | 0.177 ± 0.006 | 0.180 ± 0.006# |
| Gonadal fat pad weight, g | 1.043 ± 0.068 | 0.844 ± 0.074 | 1.066 ± 0.119 | 0.628 ± 0.048 |
| Spleen weight, mg | 0.110 ± 0.011 | 0.085 ± 0.005 | 0.116 ± 0.017 | 0.089 ± 0.006 |
| Blood glucose, mg/dl | 159 ± 5 | 150 ± 6 | 135 ± 6* | 143 ± 6 |

Values shown are the means ± SEM. Statistical analysis was performed by one-way ANOVA followed by Tukeys' multiple comparison post-test. *$p<0.05$ vs. A-SED; #$p<0.05$ vs. O-SED

Supplementary Table 2. Primers used for qPCR

| Gene | Forward | Reverse |
|---|---|---|
| *18S* | GTAACCCGTTGAACCCCATT | CCATCCAATCGGTAGTAGCG |
| *Fbn1* | GGACGCCAATTTGGAGGCT | CTTTCAGCGCATCGTGTCCT |
| *Fbn2* | CTCCACCAAAGACGCTCTGG | CCCTCGTCCCGATACTCAGG |
| *Tgfb1* | CTCCCGTGGCTTCTAGTGC | GCCTTAGTTTGGACAGGATCTG |
| *Tgfb2* | CTTCGACGTGACAGACGCT | GCAGGGGCAGTGTAAACTTATT |
| *Ctgf* | GGGCCTCTTCTGCGATTTC | ATCCAGGCAAGTGCATTGGTA |
| *Sod1* | AACCAGTTGTGTTGTCAGGAC | CCACCATGTTTCTTAGAGTGAGG |
| *Sod2* | CAGACCTGCCTTACGACTATGG | CTCGGTGGCGTTGAGATTGTT |
| *Catalase* | AGCGACCAGATGAAGCAGTG | TCCGCTCTCTGTCAAAGTGTG |

Cho et al, Graphical abstract

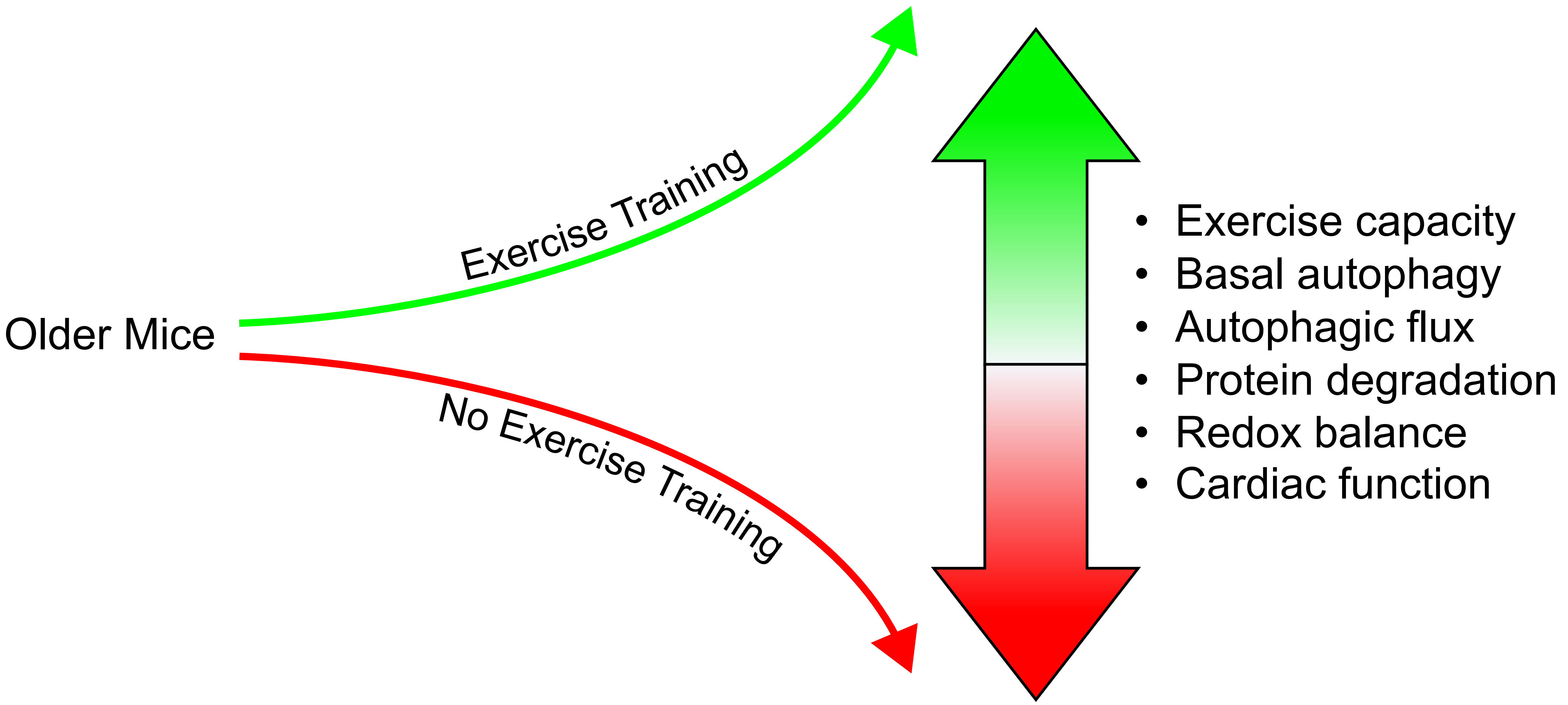
